# Activation of β2-adrenergic receptors in microglia alleviates neuropathic hypersensitivity in mice

**DOI:** 10.1101/2022.12.18.520924

**Authors:** Elisa Damo, Amit Agarwal, Manuela Simonetti

## Abstract

Drugs enhancing the availability of noradrenaline are gaining prominence in the therapy of chronic neuropathic pain. However, underlying mechanisms are not well understood, and research has thus far focused on α2-adrenergic receptors and neuronal excitability. Adrenergic receptors are also expressed on glial cells, but their roles toward antinociception are not well deciphered. This study addresses the contribution of β2-adrenergic receptors (β2-ARs) to the therapeutic modulation of neuropathic pain in mice. We report that selective activation of β2-ARs with Formoterol inhibits pro-inflammatory signaling in microglia *ex-vivo* and nerve injury-induced structural remodeling and functional activation of microglia *in vivo*. Systemic delivery of Formoterol inhibits behaviors related to neuropathic pain, such as mechanical hypersensitivity, cold allodynia and the aversive component of pain, and reverses chronically established neuropathic pain. Using conditional gene targeting for microglia-specific deletion of β2-ARs, we demonstrate that the anti-allodynic effects of Formoterol are primarily mediated by microglia. Although Formoterol also reduces astrogliosis at late stages of neuropathic pain, these functions are unrelated to β2-AR signaling in microglia. Our results underline the value of developing microglial β2-AR agonists for relief from neuropathic pain and clarify mechanistic underpinnings.

**Graphical Abstract:** 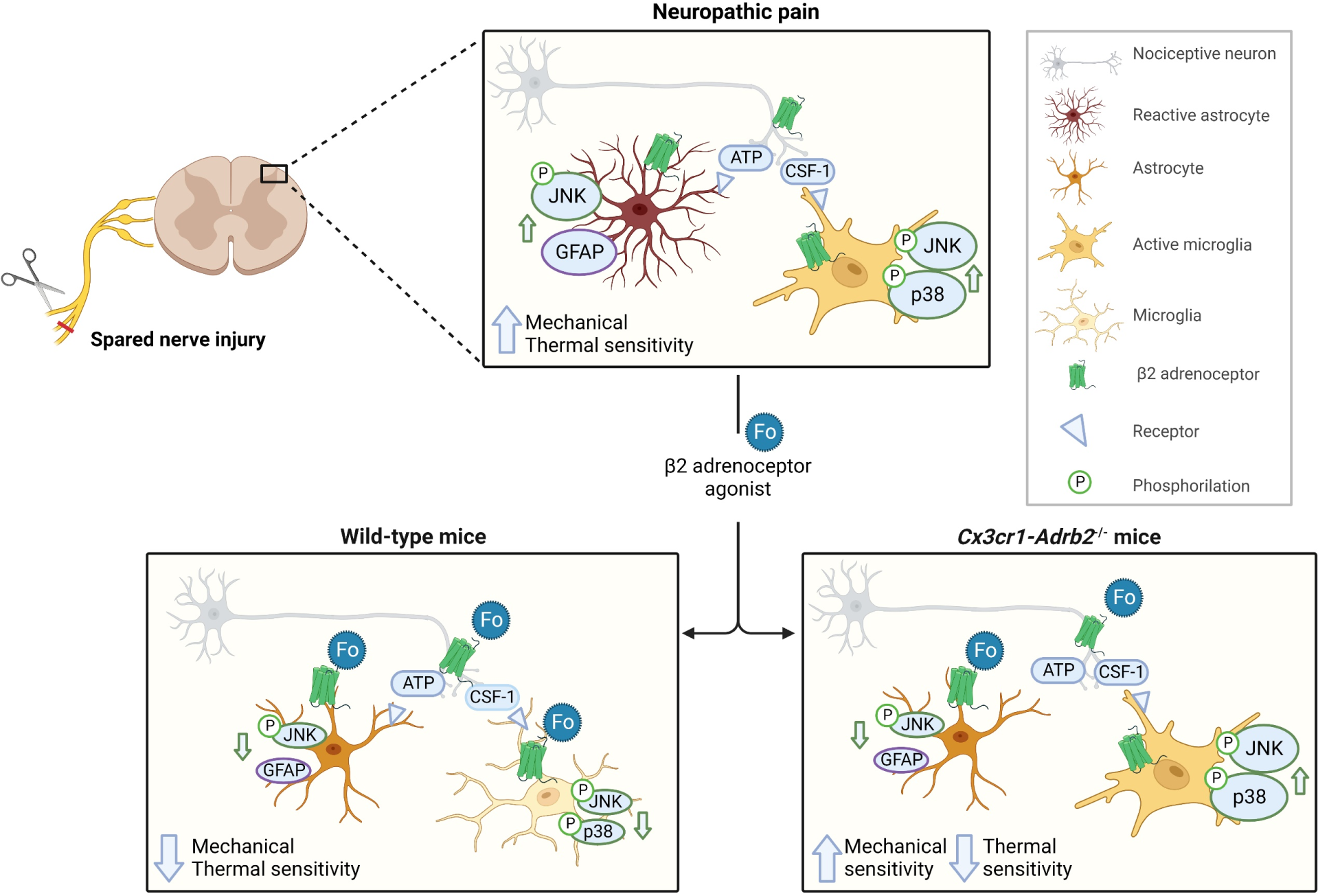

## Introduction

Chronic pain is one of the most common health problems worldwide. About seven to eight percent of the adult population suffers from chronic pain of neuropathic origin, i.e., resulting from diseases or syndromes affecting the somatosensory system or from direct nerve damage [1]. Although several analgesic drugs are available, a large proportion of patients do not respond satisfactorily to traditional treatments. Although, opioids show adequate efficacy, after a short period of treatment they lead to tolerance and strong side effects [1]. Hence, it is essential to study the molecular mechanisms of neuropathic pain to identify new molecular targets, which can be capitalized on the development of new therapies for pain relief.

Antidepressant drugs, which enhance the synaptic availability of serotonin (5-HT) and noradrenaline (NA), have emerged as key therapeutics in neuropathic pain. Several studies reported that serotonin noradrenaline reuptake inhibitors (SNRIs), a class of antidepressants, are particularly effective in treating neuropathic pain that is refractory to other treatments [1, 2], and their actions are mainly driven by enhancing NA availability [3]. NA is known to modulate pain both peripherally, where it is released by sympathetic neurons, or centrally, where it is released in the spinal dorsal horn from axons of the bulbospinal descending noradrenergic pathway originating in the locus coeruleus (LC). At the molecular level, the analgesic effect of NA has been suggested to be mediated by activation of the α2a-adrenergic receptor (α2a-AR), a Gi-coupled receptor, which on activation blocks neuronal activity [4]. Similarly, 5-HT and NA released from descending pathways promote the spinal release of enkephalins, which also exert neuronal inhibition through the activation of Gi-coupled opioid receptors [5]. In addition, activation of Gi-coupled receptors leads to the inhibition of glutamate release from primary afferent fibers as well as to the hyperpolarization of postsynaptic neurons in the dorsal horn of the spinal cord, inhibiting voltage-gated Ca^2+^ channels and opening inward rectifying K^+^ channels, respectively [6]. Activation of α2a-AR in the dorsal horn prevents neuroinflammatory changes associated with rodent models of chronic pain, reducing the release of inflammatory cytokines [7].

While efforts thus far have focused on adrenergic signaling in neurons, it is becoming increasingly evident that glial cells, including the two main central glial classes of astrocytes and microglia, express a variety of adrenergic receptors [8, 9]. In particular, astrocytes and microglia express high levels of α2a-AR and β2-AR, respectively, two receptor families known to play a role in neuropathic pain [10]. So far, however, the role of adrenergic signaling in glia in chronic pain has not been thoroughly explored.

The essential contribution of glial cells in the regulation of neural plasticity in pathways of nociception and pain is well-documented [11, 12], and glial cells have emerged to be a crucial element in the development and maintenance of chronic pain [13]. Spinally, astrocytes and microglia are proposed to be activated in a consequential manner in various models of inflammatory and neuropathic origin [12,14,15]. Interfering with their activation at an early stage can block the development of nociceptive hypersensitivity [12], whereas optogenetic activation of astrocytes in the spinal cord has been shown to be sufficient to cause nociceptive hypersensitivity [16]. Following peripheral neuronal damage, primary afferents release various substances such as colony stimulating factor 1 (CSF-1) [17] and cytokines in the spinal cord, which bind to specific receptors on microglia and activate them [18]. The activated microglia in turn produce a wide variety of active substances that act on neurons and astrocytes, initiating a crosstalk that has been suggested to sustain chronic changes in nociceptive sensitivity.

While α2-ARs have been at the center of studies addressing adrenergic modulation of pain for decades, recent studies have brought β2-ARs into focus. Systemic administration of β2-AR agonists has been shown to have anti-inflammatory and anti-nociceptive properties in long-lasting inflammatory pain [19, 20], neuropathic pain [21] and incisional pain [22]. Microglia express high levels of Gs-coupled β2-adrenoceptors (β2-AR) [23, 24] and respond to the application of NA [9,25,26]. However, given the broad distribution of β2-AR in neurons, astrocytes, microglia, and a variety of non-neuronal cells, the role of microglial β2-AR signaling remains unknown.

We designed this study to delineate the contribution of microglia to the analgesic actions of β2-AR agonists in rodent models of neuropathic pain, with emphasis on both sensory and emotional components of pain. A further goal was to study the underlying cellular and molecular mechanisms. Importantly, given the complex interplay between spinal neurons, microglia and astrocytes following nerve injury, we sought to pinpoint the temporal phases of pathophysiology induced by nerve injury in which targeting β2-AR in microglia would bring maximum benefits. Employing highly specific genetic tools, we demonstrate the key significance of adrenergic signaling in microglia via β2-AR in molecular plasticity in the spinal cord and ensuing neuropathic pain. Our findings underscore a major therapeutic potential for targeting β2-AR in neuropathic pain and yield key scientific insights into the neurobiological underpinnings of neuropathic pain.

## Materials and Methods

### Animal handling

All experimental procedures were approved by the local governing body (Regierungspräsidium Karlsruhe, Germany, Ref. 35-9185.81/G-177/17 and 35-9185.81/G-274/19) and abided by German Law that regulates animal welfare and the protection of animals used for the scientific purpose (TierSchG, TierSchVersV).

C57BL/6J mice (WT mice) of both sexes were purchased from Janvier Labs (France). Adult mice (8 weeks old, 20 – 30 g) were used for behavioral, qPCR, and immunofluorescence experiments, while 5 weeks old C57BL/6J mice were utilized for microglia primary cell culture.

To generate mice lacking β2-AR specifically in microglial cells, mice carrying a conditional allele for the *Adrb2* (*Adrb2^fl/fl^*) gene (shared by Dr. Gerald Karsenty, Columbia University, New York, USA) [27], were crossed with inducible *Cx3cr1-CreERT2* mice (shared by Dr. Steffen Jung, The Weizmann Institute of Science, Rehovot, Israel, and Dr. Frank Kirchhoff Center for Integrative Physiology and Molecular Medicine, University of Saarland, Homburg, Germany) [28] that express the tamoxifen-inducible Cre under control of microglia and macrophages specific promoter. Cre-mediated recombination of *Adrb2* floxed allele was induced in 5 weeks old *Cx3cr1-CreERT2*; *Adrb2^fl/fl^* mice by injecting intraperitoneally (i.p.) 50 mg/kg Tamoxifen (10 mg/ml, T5648 Sigma-Aldrich/Merck) once per day over 5 consecutive days. To ensure a complete loss of function of *Adrb2* protein in microglial cells, we waited for three weeks after tamoxifen treatment before proceeding with the experiments. Adult *Cx3cr1-CreERT2*; *Adrb2^−/−^*; mice (8-9 weeks old) were used for behavioral and immunohistochemistry experiments.

A total of 258 C57BL/6J mice (130 male and 128 female), 100 *Cx3cr1-CreERT2*; *Adrb2^fl/fl^* mice (50 male and 50 female) were used for the experiments.

Mice of the same sex were housed together in groups of 2 – 4 per cage and kept under a 12 h light / dark cycle at controlled temperature (22 ± 2 °C), humidity (50 – 60 %) with food and water provided ad libitum in conformity with ARRIVE guidelines.

### Surgical procedures and nerve injury

Mice of both sexes were assigned randomly and equally to spared nerve injury (SNI) or sham groups. SNI operation was performed according to earlier protocols [29] with minor modifications. Briefly, 8-9 weeks old male and female mice were anesthetized using a mix of 2 % isoflurane, oxygen, and nitrous oxide. The common peroneal and tibial nerves were exposed via an incision of the lateral thigh skin, tightly ligated, and cut distally. A 1 mm section was removed from the ligation leaving the sural nerve intact. The sham operation proceeded similarly without any nerve damage. The muscle tissue was restored, and the skin was stitched with Marlin 4-0 absorbable suture.

### Pharmacological drugs

Mice were intraperitoneally (i.p.) injected with 50μg/kg of Formoterol (cat # 1448, Tocris) [30] or its solvent (0.9 % NaCl) 1 hour before behavioral analysis or perfusion for immunofluorescence experiments.

### Behavioral tests

Animals were randomly assigned into different groups. All behavior tests were performed double-blinded, which complied with the guidelines of the International Association for the Study of Pain. The experimenter was thus unaware of the identity of the treatment groups. All behavioral measurements were done in awake, unrestrained, age and sex-matched adult mice.

#### Experimental design

Basal measurements for mechanical and cold hypersensitivity were taken twice, once per day, two days before the SNI or sham operation using von Frey filaments and cold plate test, respectively. The cold plate test was done sequentially after the von Frey filament test. Mechanical hypersensitivity and cold plate tests were performed following three different experimental plans:

1. The first one assessed the behavioral response of the mice on day three after the operation, one hour after receiving i.p. 50μg/kg of Formoterol or saline.
2. In the second paradigm, we tested mechanical and cold hypersensitivity six and 21 days after the operation, each day 1 hour after injecting i.p. Formoterol or saline (Figure 2A).
3. In the third experimental scheme, we evaluated behavioral parameters only on day 21 after the surgery, one hour after receiving i.p. Formoterol or saline (Supplementary 3A).

To analyze the mechanical and thermal (cold) response over the time after Formoterol injection, von Frey filaments measurements were taken 1, 6, 12, and 24 hours after Formoterol injection. For each time point, we tested each group of mice first with mechanical tests and then with thermal tests. Because of the technical time required to perform the mechanical test (about 2 hours), the thermal tests were performed with some delay from drug injection. This resulted in each group being tested for mechanical stimuli 1, 6, 12 and 24 after injection, while thermal tests were performed 3, 8, 14 and 26 hours after injection. To avoid undue stress to the animals, we used a different cohort of mice to test the missing time points for each test (3 hours for the mechanical test and 1 hour for the thermal test; these time points are shown in a separate graph (Supplementary 1C, D) to reduce the time interval between the different recordings and not to lose an early potential analgesic effect. The conditioned place preference (CPP) experiment was started on day 4 (Supplementary 2B) and day 32 (Supplementary 2D) after the sham or SNI operation with male and female mice as described below.

#### Mechanical sensitivity

Mice were habituated to the experimental setup, the von Frey elevated grid (Ugo Basile Inc., Italy) for 1 hour in three separate sessions within the week preceding the time of behavioral testing as well as 20 – 30 min before each testing session. Mechanical sensitivity testing was performed on an elevated grid by applying a set of von Frey filaments with increasing forces (0.008 – 1.0 g), to the affected and the contralateral hind paws using the up-down method described by Dixon [31]. Withdrawal frequencies were determined from 5 applications per filament, with a minimal interval of 5 min between filaments. Paw lifting and licking were defined as positive responses. The 50% withdrawal threshold (g) was determined by fitting the response rate vs. von Frey force curves with a Boltzmann sigmoid equation with constant bottom and top constraints equal to 0 and 100, respectively [32]. The integral of response frequency – von Frey force intensity (0.008 to 0.1 g) curves was calculated as area under the curve (AUC, A.U. = arbitrary unit).

#### Cold allodynia

Following the mechanical sensitivity test, animals were placed on a rectangular cold metal surface (4 °C, Hot/Cold Plate 35100, Ugo Basile Inc., Italy) enclosed by a Perspex cylinder, and closely monitored to record the latency of the first nociceptive response (paw lifting, shaking, licking, or jumping). A 30 s cut-off was used to prevent potential injury to the paws.

#### Conditioned place preference test

The CPP test was used to determine the ability of Formoterol in exerting pain relief from ongoing pain, as described previously [33]. Test mice were conditioned to associate one of the two compartments with pain relief. Behavioral testing was performed between 9:00 a.m. and 4:00 p.m. in mice 9 weeks of age and every session lasted 30 minutes. Baseline preferences were detected before the conditioning by placing the mice on the setup and letting them freely move between the chambers. Conditioning started one day after the baseline detection. Each day for three days, the mice were injected first with saline and after 10 min they were inserted in the assigned chamber. After at least 4 h from the saline injection, the mice were injected with Formoterol and 50 min later they were placed in the other chamber. The day after conditioning, the mice were placed on the setup and let freely move across the chambers. The CPP sessions were video-recorded and scored using Any-maze for the time spent in the two compartments. The change in time spent in the Formoterol chamber, referred as score, is calculated as the difference in time spent in the Formoterol-associated chamber on the baseline and test day.

### Immunohistochemistry, imaging, and cell counts

At three, six, and 21 days from the operation and 1 h after Formoterol or saline i.p. injection mice were perfused transcardially with cold PBS followed by 4 % PFA. The spinal column was collected and post-fixed at 4 °C overnight in 4 % PFA. The day after, the L3 – L4 spinal segment was extracted, cryopreserved overnight in 30 % sucrose, and cryosectioned at 20 µm. Sections were stored at −;20 °C.

Immunostaining was performed according to the standard protocols for immunofluorescence staining [34]. Briefly, sections were washed in PBS, incubated for 15 min in 50 mM Glycine, and blocked for 1 h in blocking solution (10% normal horse serum in PBS). The following antibodies were incubated at 4 °C overnight: rabbit-anti-Iba1 (1:500, cat # 019-19741, Wako), chicken-anti-Iba1 (1:500, cat # 234 009, Synaptic System), rabbit-anti-p-p38 (1:300, cat # 9212, Cell Signaling Technology), mouse-anti-p-JNK (1:200, cat # 9255, Cell Signaling Technology), rabbit-anti-p-JNK (1:100, cat # 4668, Cell Signaling Technology), guinea pig-anti-GFAP (1:1000, cat # 173 004, Synaptic System). Negative controls were incubated overnight in blocking solution. On the following day, sections were washed for 15 min in blocking solution and 15 min in PBST (PBS buffer plus TX-100). Successively, the secondary antibodies donkey anti-chicken Alexa 488-conjugated antibody (1:1000, cat #: A78948 ThermoFisher Scientific Invitrogen, USA), donkey anti-mouse Alexa 488-conjugated antibody (1:1000, cat #A32766 ThermoFisher Scientific Invitrogen, USA), donkey anti-rabbit Alexa 594-conjugated antibody (1:1000, cat #: A21207, ThermoFisher Scientific Invitrogen, USA), and goat anti-guinea pig Alexa 647-conjugated antibody (1:1000, cat #: A11076 Invitrogen, USA) diluted in blocking solution were incubated for 1h. Sections were washed three times for 10 min in blocking solution, treated with Hoechst 33342 (cat #: H3570 ThermoFisher Scientific) diluted 1:10000 in PBS for 15 min, and rinsed two times for 10 min in PBST. Lastly, the sections were incubated for 10 min in 10 mM TRIS/HCl before mounting them on glass slides with Mowiol (cat #: 0713.1 Carl Roth) and stored at 4°C.

For p-JNK detection, slides were incubated for 20 min into an antigen retrieval buffer (10 mM of sodium citrate, 0.05 % Tween 20, pH 6) at 80 °C in the water bath. After cooling for 30 min at room temperature. Sections were washed once in 1x PBS for 5 min followed by the standard protocol. Labeled slices were imaged with a confocal laser-scanning microscope (20x, 40x objectives: Leica TCS SP8 AOBS, Germany) using identical illumination exposure parameters for all animals. Sequential line scans were used for spinal cord sections. A montage of confocal image stacks was acquired over a depth of 12 µm. For morphological analysis of microglia, slices were imaged with an epifluorescence microscope (40x objective: Nikon Y-TV55, Germany) using identical illumination exposure parameters for all groups. Images were taken with a total depth of 5 µm. The maximum z-projection spinal cord images were applied to be evaluated via Fiji-Image J software (version 1.52p, National Institutes of Health, USA).

Counting was done in the contra and ipsilateral spinal dorsal horn (SDH), lamina I-II-III on 3-6 sections on each side across the entire region of interest. In these regions, we count the number of Iba1-positive cells and double-positive p-p38/Iba1, p-JNK/Iba1, and p-JNK/GFAP cells. FIJI-Image J cell counter plug-in was used. For fluorescence intensity detection of GFAP, the region of interest was drawn and calculated as the mean grey value. The results were then divided by the area to have the intensity density per area unit. Data are expressed as the ratio between the ipsilateral and contralateral sides. For morphological analysis, 15 – 20 cells were analyzed per mouse. The microglial soma perimeter and the process length were assessed employing FIJI-image J.

### Microglia isolation

For cell culture experiments, the entire spinal cords of 5 weeks old mice were processed for RNA extraction, and the L3 – L5 lumbar part of SDH of three 8 weeks old mice was employed and pulled together for each condition. Microglia isolation was performed using the gentle MACS Dissociator (Miltenyi Biotec). Briefly, spinal cord tissues were homogenized using the Adult Brain Dissociation Kit (mouse and rat, cat # 130-107-677, Miltenyi Biotec) and running the gentleMACS program 37C_ABDK_02 following the manufacturer instructions.

The resulting cell suspension, when used for RNA extraction, was subjected to treatment with the Myelin Removal Beads II (cat # 130-096-731, Miltenyi Biotec) adjusting the volume of the buffer and the beads according to the amount of initial tissue, whereas when used for the cell culture this step was avoided. Subsequently, the cells were incubated with CD11b Microbeads (1:10, cat # 130-049-601, Miltenyi Biotec) and microglia were isolated using a magnetic MACS separator. Sorted microglia were harvested for further cell culture or stored at −80 °C for quantitative PCR. Purity check was obtained by RT-PCR investigating specific gene markers for neurons (*Syt1 V2*), astrocytes (*Aqp4*), oligodendrocytes (*Mbp*), and microglia (*Cx3cr1*) (Table 1). Primer efficiency: 1.96 (*Cx3cr1)*, 1.98 *(Aqp4),* 2.01 *(Syt1 V2),* 2.07 (*Mbp*).

**Table 1.**
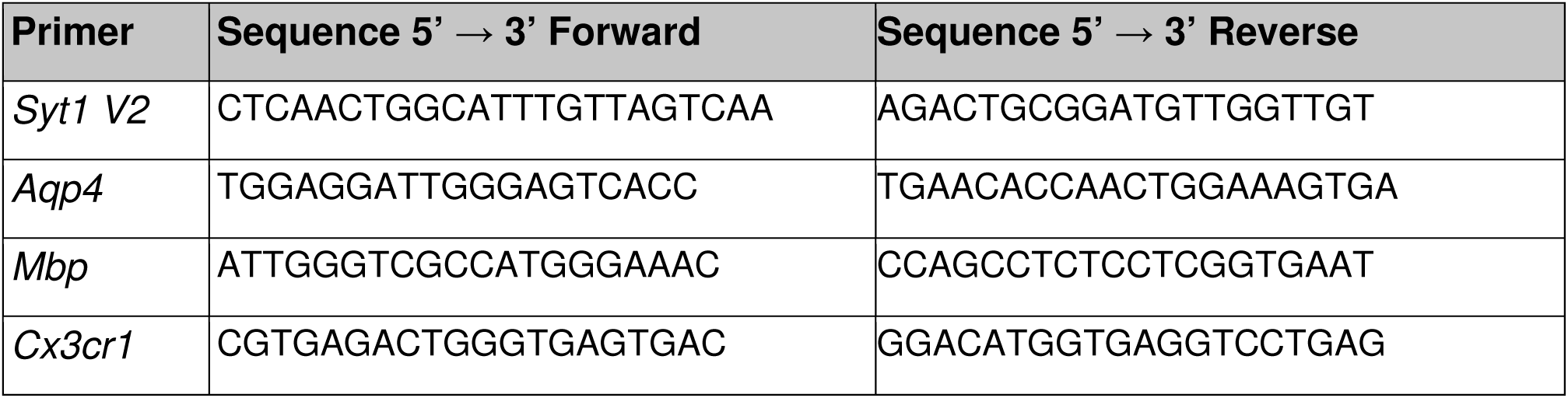
Primers used for RT-PCR to check the purity of primary microglia cell culture.

### Cell culture

Sorted microglia were resuspended in cell culture medium, plated on poly-D-Lysine (P6407-5MG Sigma Aldrich/Merck) coated glass coverslip in a 24 well/plate, and grown at 37 °C under 5 % CO2 in the incubator. Cell culture medium consists of DMEM medium (cat #: 11995-065, ThermoFisher Scientific) supplemented with Penicillin-Streptomycin (cat #: 15140-122, ThermoFisher Scientific), G-5 Supplement (1:100, cat # 17503012, ThermoFisher Scientific), IL-34 (100 ng/ml, cat # 200-34, Peprotech), cholesterol (1.5 g/ml, cat # C3045, Sigma Aldrich/Merck), TGF-β2 (2 ng/ml, cat # 100-35B, Peprotech), following the protocol from Bohlen et al., [35]. During the first five days in culture, CSF-1 (10 ng/ml, cat # 300-25, Peprotech) was added to promote cell survival and proliferation. Since CSF-1 induces an activated state of microglia cells, the primary cultures were kept for another five days in culture medium without CSF-1 to let the cells acquire a resting phenotype, or with CSF-1 to mimic neuropathic condition (activated microglia).

### Dot blot

Primary microglia cells from C57Bl/6 J mice cultured in presence of CSF-1 were treated with Formoterol (10 ng/ml, cat # 1448, Tocris) or vehicle for 1 hour. The supernatant was quickly collected, frozen, and kept at −80 °C. Collected supernatants were used to analyze the released inflammatory mediators using the Mouse Inflammation Antibody Array, Membrane 40 Targets (cat # ab133999, Abcam) following the manufacturer’s instruction.

### RNA extraction and qPCR

L3 – L5 SDH tissue and microglia from the L3 – L5 SDH were quickly collected and snap-frozen on dry ice. After extracting the total RNA using the TRIzol method (cat #: 15596018), a purification treatment with Deoxyribonuclease I Amplification Grade (cat # 18068-015) was employed as per manufacturer instructions. The first-strand cDNA synthesis was retrotranscribed using 1 µg of total RNA, oligo(dT)20 primers (cat #: 18418020), random hexamer (cat #: N8080127), and SuperScript III Reverse Transcriptase (cat # 18080044) according to the manufacturer instructions. As a control reverse transcriptase was omitted. All the reagents used for RNA extraction, purification, and first strand cDNA synthesis were from ThermoFisher Scientific.

Quantitative PCRs were run using qPCRBIO SyGreen mix separate-Rox (cat #: PB20.14-51 PCRBIOSYSTEMS) and specific primers (Table 1; Table 2) or *Gapdh* as housekeeping gene (Sigma-Aldrich), on a LightCycler 96 Real-Time PCR System (Roche). The data were analyzed using the related software. The expression level of the target mRNA was normalized to the expression of *Gapdh* mRNA. The relative gene expression was quantified using the comparative ΔΔCt method. Primer efficiency: 1.96 (*Adrb2*), 1.98 (*Gapdh*).

**Table 2.**
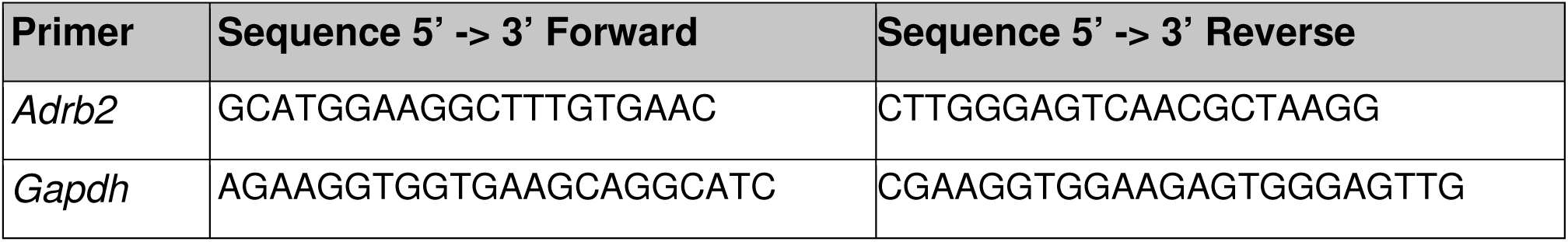
Primers used for RT-PCR to study spinal cord expression of adrenoceptors.

### Statistical analysis

All data are expressed as mean ± standard error of the mean (S.E.M). Statistical analysis was performed using Prism 9 (GraphPad Software). When compared two groups of data two-tailed, unpaired Student’s t-test was used, whereas when multiple groups and variables were compared, two-way ANOVA was employed and post-hoc Tukey’s test for multiple comparisons was performed to determine statistically significant differences. A p-value of < 0.05 was considered significant. Sample number (n), p-values, and interactions (when utilized the two-way ANOVA test) are indicated in the figure legends.

### Data availability

The authors agree to make all raw data available and will upload raw data to a repository of Heidelberg University which is currently under construction. Furthermore, the raw data of this study will be made available on a request to MS.

## Results

### β2-ARs are upregulated in spinal microglia early after nerve injury and their activation attenuates inflammatory mediators in microglia

Because activation of the noradrenergic descending pathway has been suggested to regulate neuroimmune processes, we addressed whether microglia are affected and whether β2-ARs play a role. Therefore, we isolated microglia from segment L3 – L5 of the SDH of wild-type mice using the magnetic-assisted cell sorting (MACS) technique. Next, we ascertained the purity of microglia extraction through quantitative RT-PCR (qPCR) analysis, which revealed enrichment of microglial gene and a lack of expression of genes expressed in neurons (*Syt*1 V2), astrocytes (*Aqp4*), and oligodendrocytes (*MBP*) (Figure 1A). To study whether nerve injury changes the expression of the murine gene encoding β2-ARs (*Adrb2*), we employed the spared nerve injury (SNI) model of neuropathic pain or sham surgery as control. Three days post-surgery, we extracted mRNA from MACS-sorted microglia or total spinal cord lysates from L3-L5 of SDH. Using qPCR, we examined the expression of *Adrb2* in SNI and sham conditions, while *Adrb2* expression did not differ significantly in bulk mRNA isolated from SDH samples (Supplementary 1A), we observed a significant and robust upregulation of *Adrb2* in microglia isolated from L3-L5 of SDH from mice with SNI compared to sham-operated mice (Figure 1B). Thus, we conclude that the expression of *Adrb2* is specifically upregulated in spinal cord microglia early after nerve injury.

**Figure 1.**
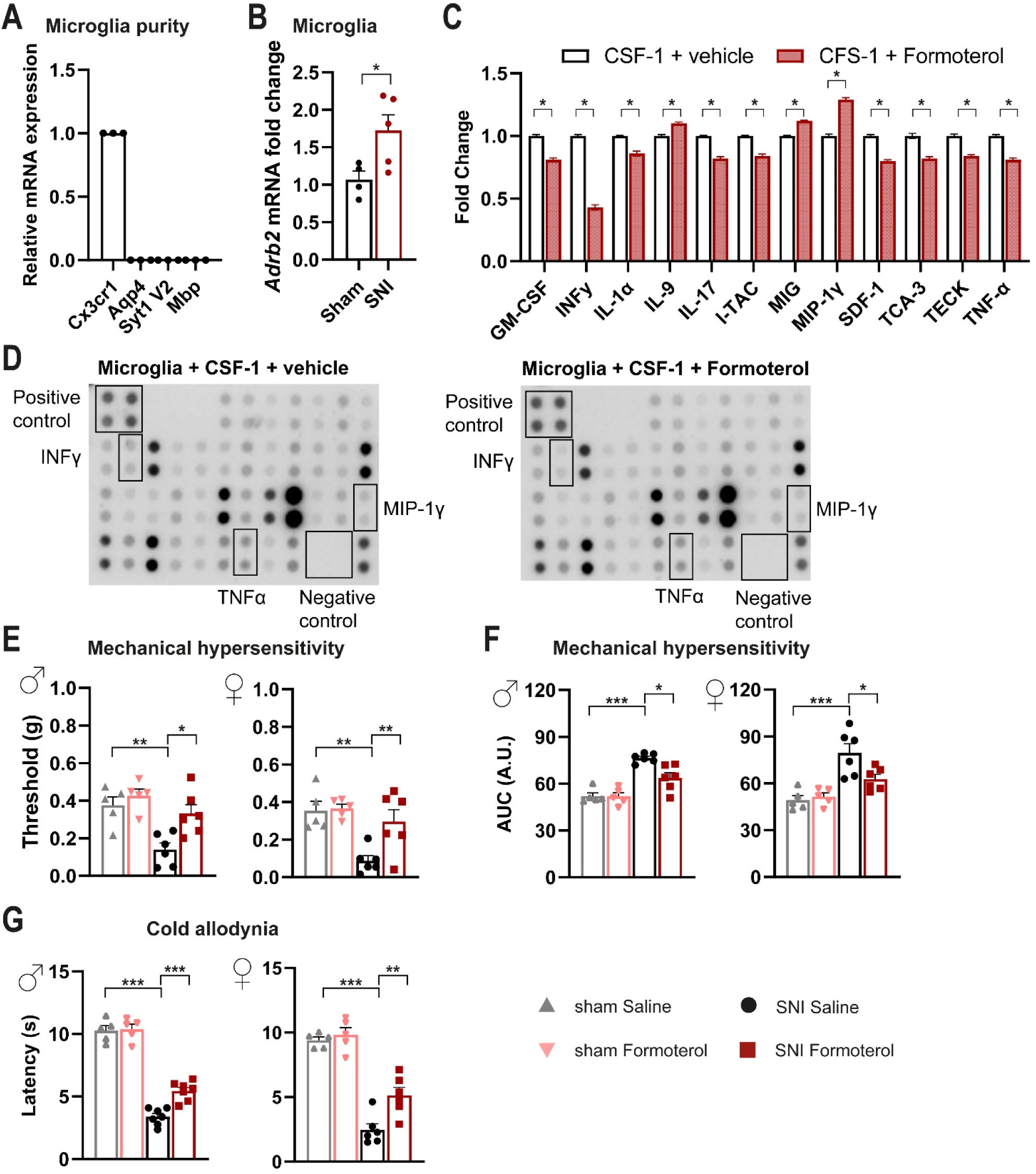
Spared nerve injury-induced induces upregulation of the *Adrb2*mRNA in microglia, whereas β2-AR agonist decreases inflammation markers in activated primary microglia cell culture and sensitization in vivo. (A) Purity check of microglia isolated from the spinal dorsal horn via qPCR on markers specific for microglia (*Cx3cr1*), astrocytes (*Aqp4*), neurons (*Syt1 V2*), and oligodendrocytes (*Mbp*). n = 3. (B) Relative expression of *Adrb2* mRNA in microglia isolated from the spinal dorsal horn of SNI- and sham-operated mice three days after surgery. n = 4 – 5 / group; two-tailed unpaired t-test was performed; * p < 0.05 as compared between two groups. (C) Activated primary microglia culture (CSF-1 treated) decreases the release of inflammatory mediators after Formoterol treatment. n = 4. (D) Examples of dot blots for inflammatory cytokines released from activated primary microglia culture after vehicle or Formoterol treatment. (E, F) Behavioral analysis of the effects of intraperitoneal Formoterol administration on mechanical sensitivity measured 1 h after intraperitoneal injection of Formoterol (E) in male (left, F1,18 = 3,835, p = 0,0659) and female (right, F1,18 = 6,184, p = 0,0229) mice. Integral of response frequency – von Frey force intensity (0.008 to 0.1 g) curves (AUC, A.U. = arbitrary unit) (F) three days after SNI or sham operation, using male (left; F1,20 = 3,169; p = 0,0903) and female (right; F1,18 = 5,561, p = 0,0299) mice. (G) Cold allodynia measured after Formoterol injection, three days after operation, using mice of both genders (male: F1,20 = 8,808, p = 0,0076; female: F1,18 = 4,913, p = 0,0398). n = 5 – 7 / group; two-way ANOVA test; * p < 0.05, ** p < 0.01, *** p < 0.001. Data are expressed as mean ± SEM, individual data points are displayed.

Furthermore, we generated primary microglia cultures from spinal cord tissue of wild-type mice. To emulate microglial activation post-nerve injury, we incubated cultured microglia with CSF-1 [17] and tested the impact of treatment with the β2-AR specific agonist Formoterol or the corresponding vehicle. We found that Formoterol treatment attenuated the CSF-1-induced expression of inflammatory mediators such as Interleukin 1 alpha (IL-1α), Interferon gamma (INF-γ), Tumor necrosis factor alpha (TNF-α), and Interleukin 17 (IL-17) (Figure 1C, D), compared to vehicle. Moreover, the levels of chemokines that induce proliferation and/or promote chemotaxis of immune cells to the injury site (Granulocyte-macrophage colony-stimulating factor (GM-CSF), Interferon-inducible T cell alpha chemoattractant (I-TAC), Stromal cell-derived factor 1 (SDF-1), Chemokine ligand 1 (TCA-3/CCL1) and Chemokine ligand 25 (TECK/CCL25)) were also decreased. Expression of anti-inflammatory cytokines such as Interleukin 9 (IL-9) increased in the Formoterol-treated cultures, as well as other chemotactic cytokines for migrating immune cells as Macrophage Inflammatory Protein 1 Gamma (MIP-1γ/CCL9) and Monokine induced by interferon-gamma (MIG) (Figure 1C, D). Thus, activation of β2-ARs in microglia suppresses pro-inflammatory signaling and response of microglia.

### Impact of in vivo administration of β2-AR agonist on mechanical and cold hypersensitivity in mice over early stages post-nerve injury

Formoterol has been previously reported to exert an anti-nociceptive effect in a neuropathic pain model [30], but its mechanism is yet not clarified. We chose to apply Formoterol systemically via intraperitoneal (i.p.) delivery to facilitate therapeutic relevance and studied its anti-nociceptive effect in the SNI model of neuropathic pain. Three days post-SNI a detailed time-course analysis of behavioral responses to mechanical and cold stimuli showed that the maximum inhibitory effect on mechanical hyperalgesia is reached 1 hour (h) after Formoterol injection, whereas peak inhibition of cold hypersensitivity occurs at 3 h (Supplementary 1C, D). In both male and female mice, Formoterol injection significantly reduced SNI-induced mechanical hypersensitivity, compared to saline-treated SNI mice (Figure 1E, F; the integral of the response frequency-von Frey stimulus intensity curve from 0.008 to 1.0 g is shown). Moreover, in both genders, Formoterol application significantly increased the latency of paw withdrawal after cold stimuli compared with SNI-operated vehicle-treated mice, indicating a reduction in SNI-induced cold allodynia (i.e., when non-noxious cold is perceived as noxious) (Figure 1G). These results thus indicate that systemic delivery of a β2-AR agonist attenuates mechanical and cold hypersensitivity over the early stages of post-nerve injury.

### Formoterol reverses hypersensitivity established over several days to weeks and alleviates spontaneous pain in male mice post-SNI

Our analyses at early stages post-SNI (Figure 1E - G), uncovered the potential of systemically applied Formoterol in inhibiting the development of nociceptive hypersensitivity to mechanical and cold stimuli. Thus, we tested the impact of Formoterol in reversing hypersensitivity once it is established over several days to weeks (Figure 2A).

**Figure 2.**
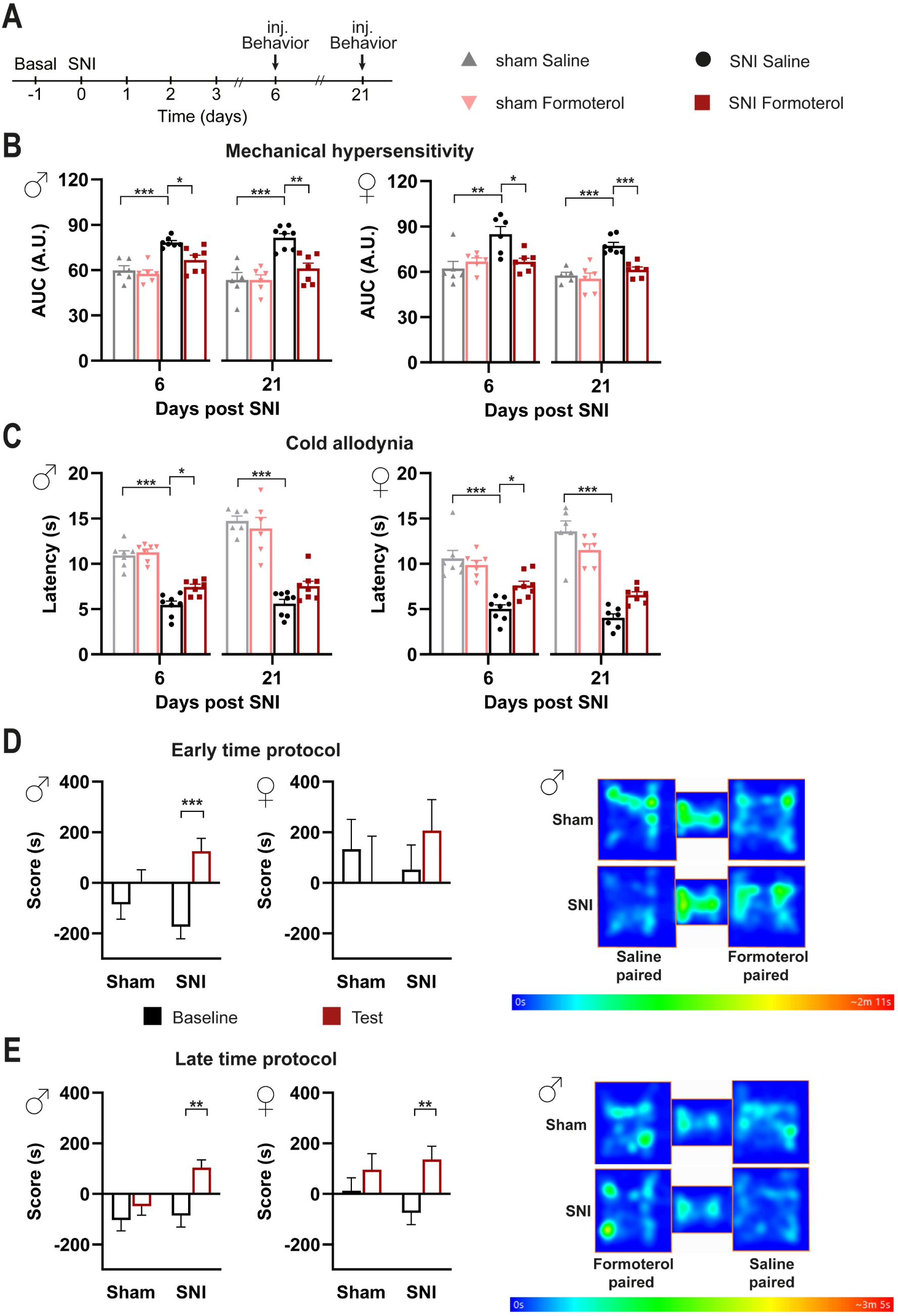
Impact of Formoterol on behavior in SNI- and sham-operated WT mice. Conditioned place preference test shows that Formoterol reduces spontaneous pain in SNI-operated mice. (A) Experimental scheme used for testing mechanical sensitivity (von Frey) and cold sensitivity (cold plate) at the plantar hind paw with 8 weeks old mice. Inj. = injection. (B) Mechanical sensitivity was evaluated as integral of the response frequency-von Frey stimulus intensity from 0.008 to 1.0 g (AUC, A.U. = arbitrary unit) six and 21 days after operation in male (left; day 6: F1,22 = 3,338, p = 0,0813; day 21: F1,23 = 8,048, p = 0,0093) and female (right; day 6: F1,21 = 9,356, p = 0,0060; day 21: F1,21 = 7,224, p = 0,0138) mice, injected with saline or Formoterol. (C) Response to cold allodynia six and 21 days post-surgery is indicated as latency (s) of paw withdrawal for males (left; day 6: F1,25 = 3,972, p = 0,0573, day 21: F1,24 = 3,770, p = 0,0640) and female (right; day 6: F1,26 = 7,740, p = 0,0099; day 21: F1,22 = 10,98, p = 0,0032) mice. n = 6 – 8 / group; two-way ANOVA test was performed. * p < 0.05, ** p < 0.01, *** p < 0.001. Data are expressed as mean ± SEM, individual data points are displayed. (D) Conditioned place preference (CPP) was tested to Formoterol at 4 days post-SNI or sham surgery. Analysis of the time spent by operated male (left) or female (right) mice is shown in the Formoterol-paired chamber before (baseline) and after (test) the conditioning. Example of heat maps recorded on the test day. (E) CPP was tested to Formoterol at 32 days post-SNI or sham operation. Score = the time spent by operated male (left) or female (right) mice In the Formoterol-paired chamber on baseline and test day. Representative example of heat maps recorded on the test day. n = 6 – 7 / sham group; n = 8 / SNI group; two-tailed unpaired t-test was performed; ** p < 0.01, *** p < 0.001 as compared between Formoterol-paired chamber baseline and test daytime. Data are indicated as mean ± SEM.

In comparison to saline application, Formoterol application at days six or 21 post-SNI led to a significant decrease in neuropathic hypersensitivity to mechanical stimuli (Figure 2B, Supplementary 2A) and cold stimuli (Figure 2C). Indeed, Formoterol injection diminished the exaggerated cumulative response to all the filaments tested (from 0.008 to 1 g) on day 6 and day 21 post-SNI (Figure 2B, Supplementary 2A). Similar observations were made with respect to allodynia to a cold stimulus on day six post-SNI (Figure 2C). In contrast, Formoterol was not able to significantly inhibit cold allodynia at day 21 post-SNI in both female and male mice (Figure 2C). These data reveal the anti-allodynic effects of systemic Formoterol at late stages after nerve injury and indicate the importance of β2-AR in mediating these effects at early and late stages of mechanical allodynia and early stage of cold allodynia.

Spontaneous pain is a key symptom in patients suffering from neuropathic [36]. To study spontaneous pain in mice, we performed the conditioned place preference (CPP) test, in which mice are conditioned to analgesic treatment, such as with pregabalin, in a chamber with specific contextual cues. Preference for the chamber on a day of testing in the absence of pregabalin is employed as a parameter indicative of ongoing pain [37] and the test is well-established in the SNI model [38]. Here, we tested whether Formoterol could induce CPP in a manner similar to pregabalin in neuropathic mice. In the CPP protocol, analgesia must be limited to the time spent in the conditioned chamber, so that the animals can unambiguously associate pain relief with the chamber in which the analgesic drug was administered. Since

Formoterol-induced antinociceptive effects last for less than 6 hours after a single application, this prerequisite was fulfilled in our experiments (Supplementary 1C, D). At 8 days post-SNI operation (schematic in Supplementary 2B), male mice spent more time in the Formoterol-paired chamber post-conditioning, while sham mice did not develop any preference (Figure 2D). In contrast, female SNI mice did not show a statistically significant difference as compared to the sham-operated group, although there was a tendency for an increased time spent in the Formoterol-paired chamber (Figure 2D and Supplementary 2C). To test the relevance of these findings to a more clinically relevant setting of patients with established neuropathic pain, we tested CPP to Formoterol at a late time point, namely 36 days post-SNI (Figure 2E and Supplementary 2D, E). Compared to the sham mice, male mice with SNI preserved the preference for the Formoterol-paired chamber (Figure 2E). Interestingly, female SNI mice also developed a preference for the Formoterol chamber at this late stage post-SNI (Figure 2E and Supplementary 2E). These findings suggest that Formoterol can reduce ongoing pain in both male and female mice at late stages post-nerve injury.

### In vivo administration of Formoterol dampens structural remodeling and activation of microglia in neuropathic mice

Given their role as resident immune cells in the central nervous system (CNS), microglial cells are highly sensitive to inflammatory mediators. In neuropathic pain conditions, several cell types produce a variety of pro-inflammatory substances, many of which can consequentially push microglia toward an inflammatory state [15]. We studied the impact of β2-AR activation on spinal cord microglia three, six, or 21 days after SNI by studying microglial density and morphological changes. Microglia density was analyzed as the density of Iba1-positive cells in the lamina I - III of the SDH (Figure 3A, C). Additionally, microgliosis is differentiated by shifts from ramified-homeostatic microglia towards ameboid phenotype (enlargement of soma and shortening of processes length). To describe the reactive state of microglia, we studied three hallmarks of microgliosis: density, soma perimeter, and process length (Figure 3). Since it is well known that there are sex differences in microglial contribution to the development and maintenance of neuropathic pain [39], we parsed our analysis based on the sex of mice. We evaluated the ratio between ipsilateral and contralateral SDH, in SNI-operated mice treated with saline or Formoterol. Male mice demonstrated a significant increase in microglial density in the superficial layers of the ipsilateral SDH (marked in dashed lines in examples shown in Figure 3A) at both early and late stages post-SNI (Figure 3C), which was inhibited by Formoterol treatment. In female mice, increased microglial density in the superficial SDH was noted only over early stages post-SNI, which was reversed fully by Formoterol treatment (Figure 3A, B).

**Figure 3.**
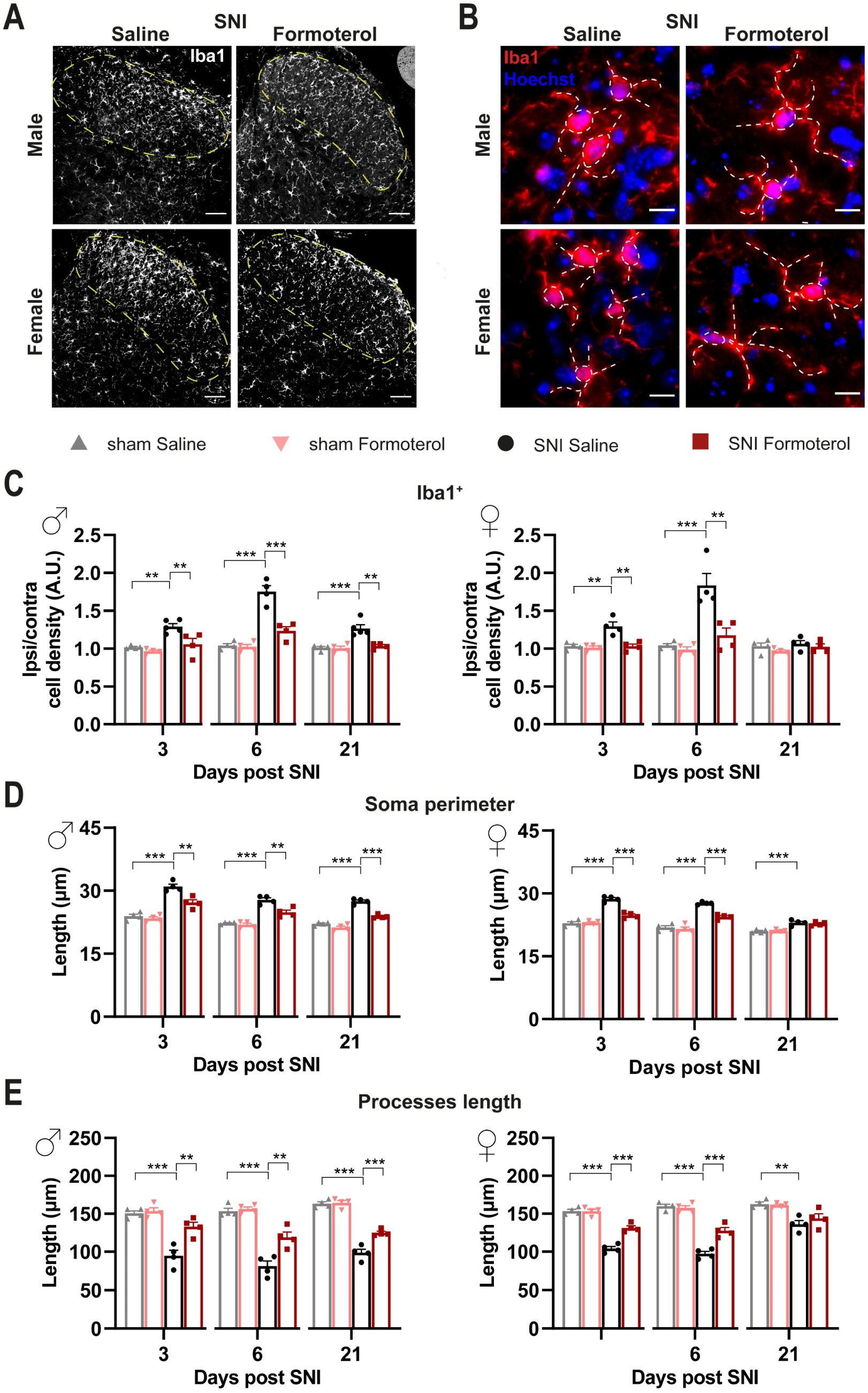
β2-AR agonist diminishes microgliosis in the spinal dorsal horn of SNI-operated mice. (A) Typical examples of Iba1-positive cells in the ipsilateral spinal dorsal horn (SDH) of male or female mice six days after the SNI surgery, injected with saline or Formoterol. Scale bar = 60 μm. (B) Representative examples of microglia (Iba1-positive with Hoechst counterstaining for cell nuclei) in the ipsilateral SDH, with somata and processes marked by white, dashed lines, six days after SNI operation injected with saline or Formoterol. Scale bar = 10 μm. (C) Analysis of cell density of Iba-1 positive microglia of male (left; day 3: F1,13 = 4,350, p = 0,0573; day 6: F1,12 = 22,62, p = 0,0005; day 21: F1,13 = 10,86, p = 0,0058) and female (right, day 3: F1,12 = 11,05, p = 0,0061; day 6: F1,12 = 10,28, p = 0,0075; day 21: F1,12 = 0,09192, p = 0,7669) treated with Formoterol or saline three, six, and 21 days after the operation. Ipsi/contra = ratio between the density value of the ipsilateral and contralateral SDH. A.U. = arbitrary unit. (D – E) Analysis of microglia morphology; the perimeter of the soma (E) and the processes length (E) in Formoterol- or saline-injected male (left; soma day 3: F1,12 = 9,207, p = 0,0104; soma day 6: F1,12 = 9,737, p = 0,0088; soma day 21: F1,12 = 22,64, p = 0,0005; processes day 3: F1,12 = 10,89, p = 0,0063; processes day 6: F1,12 = 10,78, p = 0,0065; processes day 21: F1,12 = 14,27, p = 0,0026) and female (right, soma day 3: F1,12 = 37,65, p < 0,0001; soma day 6: F1,12 = 16,83, p = 0,0015; soma day 21: F1,12 = 0,4001, p = 0,5389; processes day 3: F1,12 = 26,86, p = 0,0002; processes day 6: F1,12 = 20,92, p = 0,0006; processes day 21: F1,12 = 1,159, p = 0,3028) mice three, six, or 21 days after surgery. n = 4 – 5 / group; two-way ANOVA test was performed; ** p < 0.01, *** p < 0.001. Data are shown as mean ± SEM, individual data points are displayed.

In terms of morphological changes in spinal cord microglia in response to nerve injury, we observed that both male and female mice showed a significant increase in the perimeter of microglial cell-body with a corresponding significant decrease in the length of microglia processes over three, six and 21 days post-SNI, although the late stage changes were much more pronounced in male mice as compared to female mice (examples in Figure 3, quantitative summary in Figure 3B, D, E and negative controls in Supplementary 4C). Formoterol partially, but significantly, reversed SNI-induced microglial enlargement of the soma and the shrinking of the processes in male mice at all time points, while it failed to reverse microglial changes at late time points in female mice (Figure 3B, D, E). Interestingly, Formoterol application has no effect on microglia density or morphology in sham-operated mice (Supplementary 4A, B).

An increase in microglial density does not necessarily correspond to a reactive state of microglia. Hence, to investigate the effect of Formoterol treatment on microglia activation in neuropathic conditions, we performed co-localization analysis using Iba1 as a microglial marker and the active (i.e., phosphorylated) form of two different markers for activation: phosphorylated versions of the MAPKs p38 and JNK. Microglia are the main p38-expressing cells of the spinal cord [40]. JNK signaling modulates apoptosis and the level of pro-inflammatory cytokines, while p38 is involved in the development and maintenance of neuropathic nociceptive hypersensitivity via induction of inflammatory mediators [3]. Immunoreactivity for phosphorylated p38 in Iba1-expressing microglia was significantly enhanced in male and female mice at early (3 days post-SNI) and late (21 days post-SNI) time points after SNI and was completely reversed by Formoterol treatment at all time points (Figure 4A, B). Phosphorylated JNK was enhanced in microglia of male and female mice at three, six and 21 days post-SNI and was fully reversed by Formoterol in both cases except for female mice at 21 days after SNI operation (Figure 4C, D).

**Figure 4.**
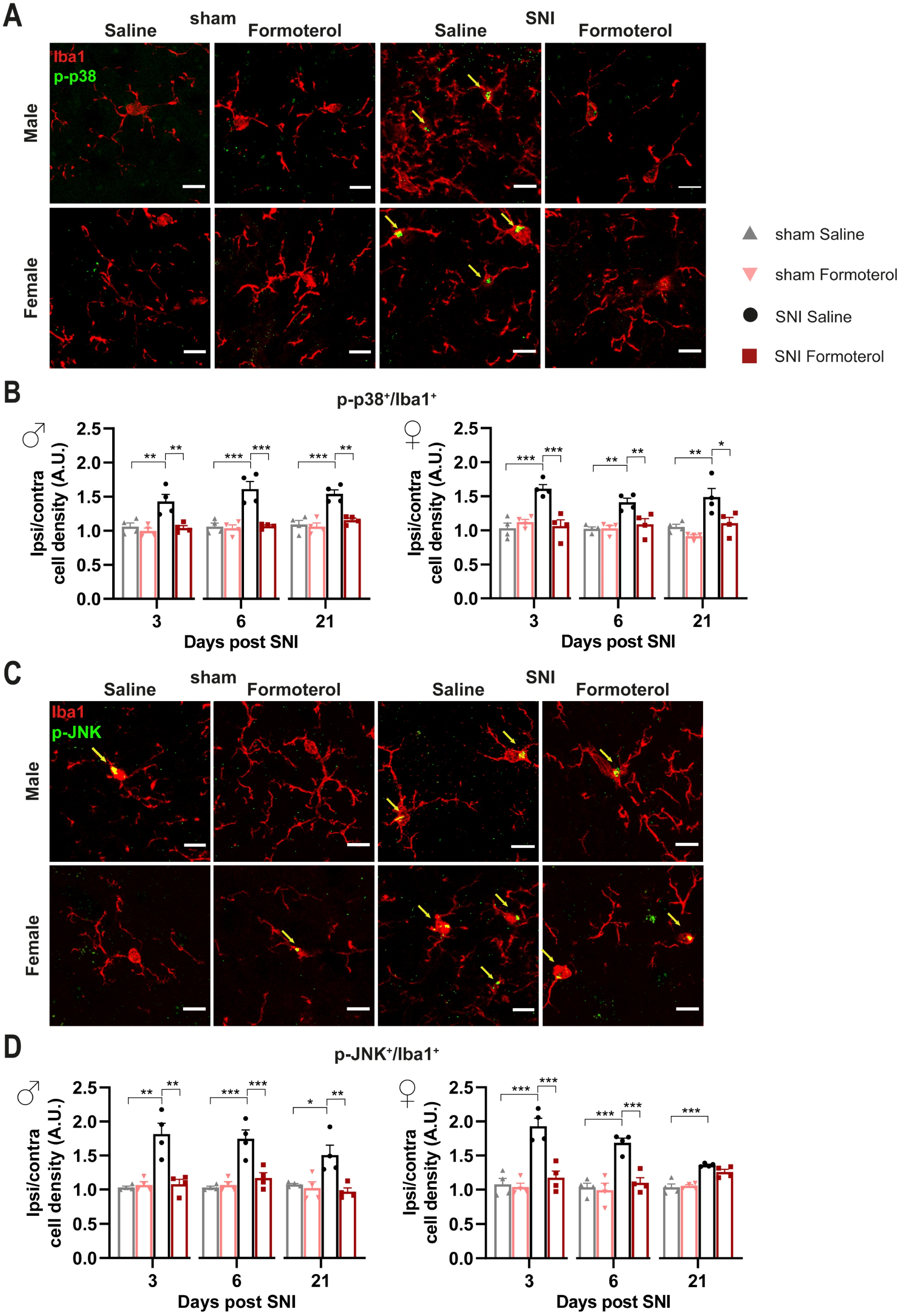
Formoterol weakens microglial activation markers in the spinal dorsal horn of SNI-operated mice. (A – B) Representative examples of colocalization of Iba1-positive signal and the microglial activation markers, p-p38 (A) and p-JNK (B), in the spinal dorsal horn in male and female mice, six days after operation injected with saline or Formoterol. Double-positive cells are pointed by arrows. Scale bar = 10 μm. (C) Colocalization analysis of the activity marker p-p38 and Iba1 immunohistochemistry three, six, and 21 days after surgery in male (left; day 3: F1,12 = 6,529, p = 0,0252; day 6: F1,12 = 15,50, p = 0,0020; day 21: F1,12 = 11,09, p = 0,0060) and female (right; day 3: F1,12 = 12,52, p = 0,0041; day 6: F1,12 = 8,325, p = 0,0137; day 21: F1,12 = 11,01, p = 0,0061) mice. (D) Analysis of the colocalization of Iba1-positive signal and the activation marker p-JNK in the spinal dorsal horn of male (left; day 3: F1,12 = 9,548, p = 0,0094; day 6: F1,12 = 15,84, p = 0,0018; day 21: F1,12 = 7,225, p = 0,0197) and female (right; day 3: F1,12 = 15,09, p = 0,0022; day 6: F1,12 = 12,27, p = 0,0044; day 21: F1,12 = 3,269, p = 0,0957) mice treated with saline or Formoterol, three, six, and 21 days after the operation. Ipsi/contra = ratio between the ipsilateral and contralateral dorsal horn of the spinal cord. A.U. = arbitrary unit. n = 4 / group; two-way ANOVA test was performed; * p < 0.05, ** p < 0.01, *** p < 0.001. Data are shown as mean ± SEM, individual data points are displayed.

Taken together, analysis of reactive microgliosis after Formoterol treatment in neuropathic mice suggests that Formoterol treatment restores normal microglial morphology and function largely over the development of neuropathic pain in both sexes. Although there was some variability, our analysis suggested that microglial activation is reversed by Formoterol in late phase after neuropathic pain induction (21 days post-SNI) in male mice, but not in female mice.

### Formoterol diminishes astrocytic activation at late stages after nerve injury in female mice

Astrocytes also express β2-AR, therefore Formoterol might exert its analgesic effect through the activation of this receptor on astrocytes. Using the same experimental protocol, we studied the effect of β2-AR activation on astrocytes using Glial fibrillary acidic protein (GFAP) to identify astrocytes in combination with a classic activation marker, p-JNK. In both sexes, we observed a significant increase in GFAP signal intensity at late time points after nerve injury (21d post-SNI) when neuropathic pain is fully developed, but not over early time points (Figure 5A, B). Interestingly SNI-induced astrogliosis was partially, but significantly, inhibited by Formoterol in both sexes (Figure 5A, B).

**Figure 5.**
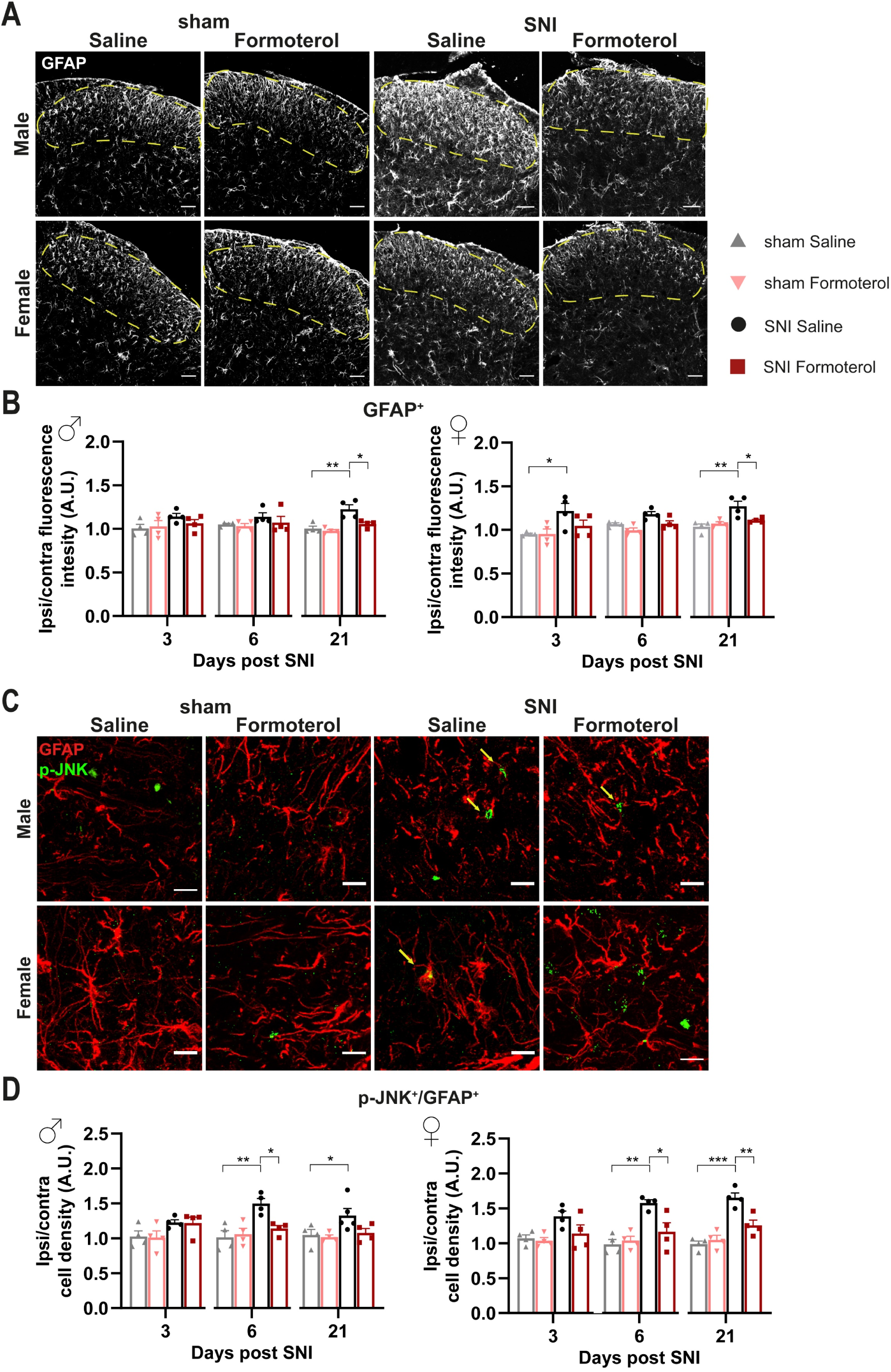
Spinal astroglial cells response to Formoterol in SNI-operated wild-type mice. (A) Examples of GFAP immune reactivity in the ipsilateral spinal dorsal horn (SDH) 21 days after nerve injury in male and female WT mice who received saline or Formoterol injection. Scale bar = 60 μm. (B) Typical examples of colocalization of GFAP astrocytic marker with glial activation marker, p-JNK in the SDH of male and female mice, 21 days after sham or SNI operation in mice injected with saline or Formoterol. Scale bar = 10 μm. (C) Analysis of GFAP fluorescent intensity of male (left; day 3: F1,12 = 1,097, p = 0,3156; day 6: F1,12 = 0,9393, p = 0,3516; day 21: F1,12 = 5,003, p = 0,0451) and female (right; day 3: F1,12 = 1,835, p = 0,2005; day 6: F1,12 = 0,4848, p = 0,4995; day 21: F1,12 = 8,059, p = 0,0149) WT mice treated with saline or Formoterol, three, six, and 21 days after the operation. Ipsi/contra fluorescent intensity = ratio between values of fluorescent intensity obtained from the ipsilateral and contralateral SDH. (D) Quantitative analysis of the colocalization of GFAP and the activation marker p-JNK in the SDH of male (left; day 3: F1,12 = 0,02015, p = 0,8895; day 6: F1,12 = 7,089, p = 0,0207; day 21: F1,13 = 2,850, p = 0,1152) and female (right; day 3: F1,12 = 1,759, p = 0,2095; day 6: F1,12 = 8,117, p= 0,0146; day 21: F1,12 = 12,89, p = 0,0037) mice treated with saline, or Formoterol, three, six, and 21 days after sham or SNI operation. Ipsi/contra = ratio between the ipsilateral and contralateral SDH. A.U. = arbitrary unit. n = 4 / group; two-way ANOVA test. * p < 0.05, ** p < 0.01, *** p < 0.001. Data are indicated as mean ± SEM, individual data points are displayed.

These findings are in line with the emerging view that microglia are activated during an acute phase and drive neuroinflammation, which lead to transition of acute pain into chronic pain, while spinal cord astrocytes contribute to central sensitization and the maintenance of chronic pain [39]. Interestingly, nerve injury also induced upregulation of p-JNK in GFAP-positive astrocytes in female mice at six and 21 days post-SNI, which was reversed by Formoterol (Figure 5C, D). Neither p-JNK upregulation in astrocytes nor effects of Formoterol were reliably seen in male and female mice at early time points post-SNI (Figure 5D).

These results suggest a disconnection between the role of β2-ARs in early microglial changes and late astrogliosis. However, there is ample literature suggesting that early microglial activation is linked to late astrocytic involvement in neuropathic pain. In our analyses, because Formoterol was sequentially administered at six and 21 days, it could not be ruled out that the Formoterol effects seen at 21 days (mostly involving astrocytes) were aided by injection of Formoterol at six days post-SNI that reduces the microglial pro-inflammatory response. To dissect these from one another, we performed an additional experiment in which Formoterol was only injected at 21 days post-SNI in wild-type mice (scheme shown in Supplementary 3A). Interestingly, in both sexes, the single late injection of Formoterol significantly reduced mechanical hypersensitivity (Supplementary 3B), and the magnitude of this change was comparable to that from our previous experiment in which Formoterol had been injected on six and 21 days post-SNI. Similar results were obtained for the cold allodynia test (Supplementary 3C). Microglial density in the SDH was not significantly changed by the single late application of Formoterol in mice of both sexes (Supplementary 3D, E). Finally, astrogliosis seen at the late time point post-SNI was significantly reduced by Formoterol given as a single application on day 21 post-SNI (Supplementary 3F, G). Taken together, these results demonstrate that Formoterol effects at late time points are not dependent on the application of Formoterol at early stages after nerve injury and suggest a disconnection between β2-AR modulation in microglia and astrocytes.

### Contribution of microglial *β*2-ARs to anti-nociceptive effects of Formoterol in mice with neuropathic pain

Our analyses uncovered the potential of systemically applied Formoterol in inhibiting microglia activation in neuropathic pain condition. Therefore, we addressed the contributions of microglial β2-ARs to Formoterol-induced analgesia, given their broad expression across cell types. To test the importance of the microglial β2-AR, we generated a conditional knockout mouse line in which the *Adrb2* gene was specifically deleted from microglia (strategy shown schematically in Figure 6A). We crossbred *Cx3cr1*-CreERT2 mice with the *Adrb2* floxed mice to generate double transgenic mice *Cx3cr1-CreERT2; Adrb2^fl/fl^* mice. *Cx3cr1-Adrb2^fl/fl^* and control (*Adrb2fl/fl*) mice were injected with tamoxifen at 5 weeks of age, and post 3-4 weeks qPCR analysis on MAC sorted microglial cells showed more than 80% reduction in the expression of Adrb2 in *Cx3cr1-Adrb2^fl/fl^* mice when compared to control mice or *Cx3cr1-Adrb2^fl/fl^* mice not injected with tamoxifen (Figure 6B).

**Figure 6.**
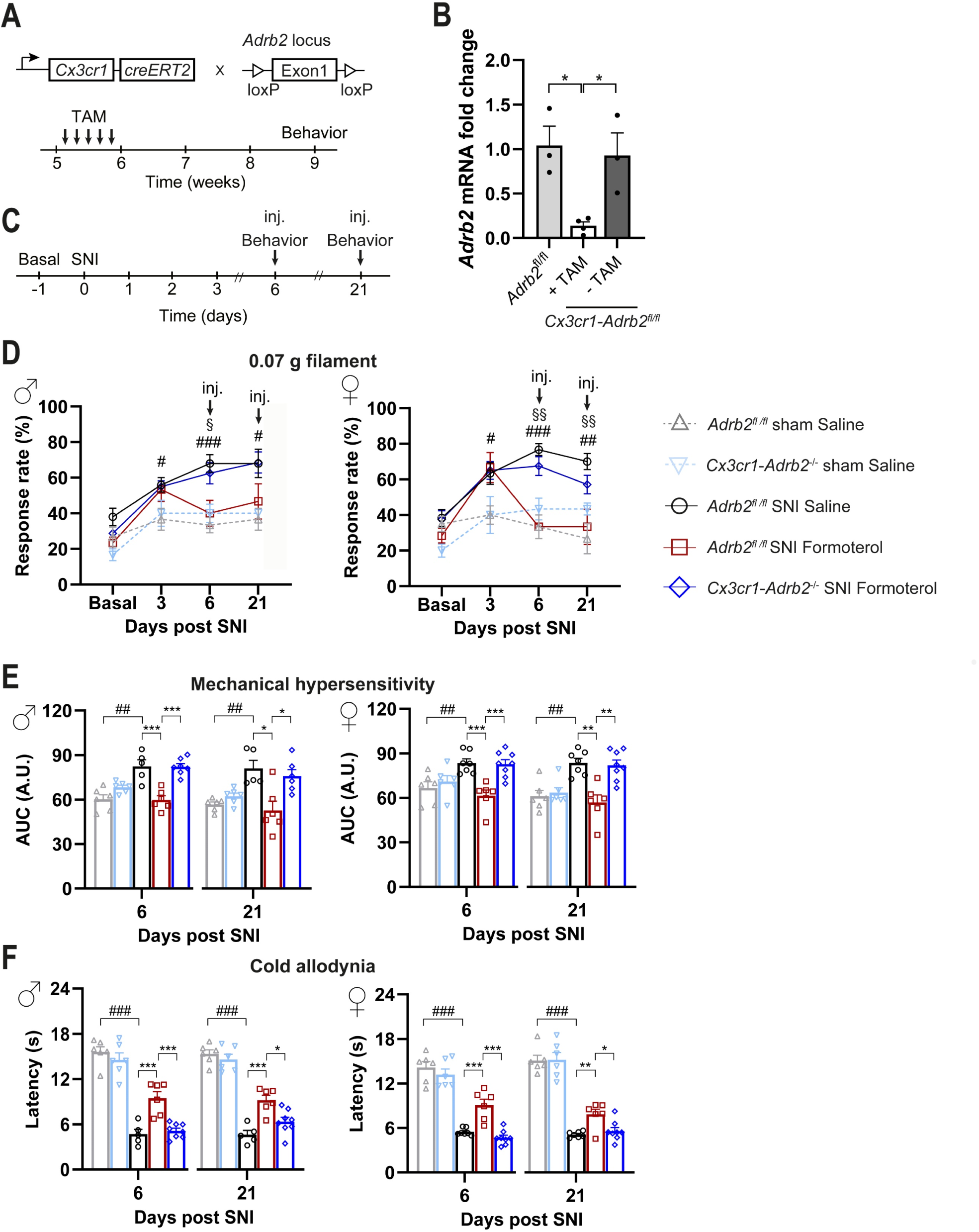
Effect of Formoterol on behavior in *Cx3cr1-Adrb2^−/−^* operated mice. (A) Schematic representation of the strategy for generation of mice lacking the *Adrb2* gene conditionally in microglial cells in a tamoxifen-inducible manner. (B) Analysis via qPCR demonstrating the loss of *Adrb2* mRNA expression in microglia cells three weeks after tamoxifen injection in the mouse line *Cx3cr1-Adrb2^fl^*^/^*^fl^*. The recombination does not occur in mice lacking the Cre-cassette (*Adrb2^fl/fl^*). TAM = Tamoxifen. n = 3 – 4 / group; ordinary two-way ANOVA with main effects only was performed. * p < 0.05. (C) Experimental scheme for testing mechanical sensitivity (von Frey) and cold allodynia (cold plate) at the plantar hind paw with 8-9 weeks old mice. Inj. = injection. (D) Mechanical sensitivity is displayed as response frequency to the 0.07 g filament in male (left) and female (right) control mice (grey line for *Adrb2^fl/fl^* sham saline-injected mice; pink line for *Cx3cr1-Adrb2^−/−^* sham saline-injected mice black line for *Adrb2^fl/fl^* SNI saline-injected mice; red line for *Adrb2^fl/fl^* SNI Formoterol-injected mice) and *Cx3cr1-Adrb2^−/−^* transgenic mice (blue line) before the SNI operation (basal measurement), three days after surgery, and six and 21 days after the SNI, one hour after Formoterol injection. n = 5 – 8 / group; t-test test was performed; § p < 0.05, §§§ p < 0.001 as compared *Cx3cr1-Adrb2^−/−^* SNI Formoterol and *Adrb2^fl/fl^* SNI Formoterol; # p < 0.05, ### p < 0.001 as compared *Adrb2^fl/fl^* SNI Saline and *Adrb2^fl/fl^* Sham Saline. (E) Mechanical sensitivity of the same groups as (D) shown as integral of the response frequency-von Frey stimulus intensity from 0.008 to 1.0 g (AUC, A.U. = arbitrary unit). (F) Cold sensitivity six and 21 days after SNI or sham surgery, after Formoterol or saline injection showed as latency (s) of paw withdrawal. n = 5 – 8 / group; t-test test was performed; # p < 0.05, ### p < 0.001 as compared *Adrb2^fl/fl^* sham saline and *Adrb2^fl/fl^* SNI saline; ordinary two-way ANOVA with main effects only was performed among SNI *Adrb2^fl/fl^* saline, *Adrb2^fl/fl^* fl Formoterol, and *Cx3cr1-Adrb2^−/−^* Formoterol; * p < 0.05, ** p < 0.01, *** p < 0.001. Data are expressed as mean ± SEM, individual data points are exhibited.

Importantly, the deletion of β2-AR receptor specifically in microglia does not significantly affect the course of allodynia/hyperalgesia following SNI. Indeed, *Cx3cr1-Adrb2*^−/−^ mice show similar baseline sensitivity and development of hypersensitivity following SNI operation comparable with those of control mice (Figure 6D, time points Basal and 3 days after SNI). In comparison to saline application, Formoterol application at days six or 21 post-SNI led to a significant decrease in neuropathic hypersensitivity to mechanical stimuli (Figure 6D, E) and cold stimuli (Figure 6F) in control mice, but not in *Cx3cr1-Adrb2*^−/−^ mice (Figure 6D-F). Figure 6D shows response frequency to a von Frey stimulation at 0.07 g force, which at basal condition elicits almost no responses, but leads to exaggerated response frequency at six and 21 days post-SNI. However, after 1 hour from i.p. injection of Formoterol, the mechanical hypersensitivity is fully reversed to baseline levels in control mice but not in *Cx3cr1-Adrb2*^−/−^ mice, thereby, suggesting that microglial β2-ARs are required for Formoterol-induced analgesia. In *Cx3cr1-Adrb2*^−/−^ mice, Formoterol failed to attenuate the exaggerated cumulative response to all the von Frey filaments tested on control mice, on day six and day 21 post-SNI (Figure 6E). Mice of both sexes showed comparable loss of Formoterol-induced analgesia and there was no apparent sexual dimorphism. Similar observations were made with respect to allodynia to a cold stimulus on day six and day 21 post-SNI. Indeed, the analgesic effect of Formoterol related to cold stimuli was lost in *Cx3cr1-Adrb2*^−/−^ mice (Figure 6F). These data reveal anti-allodynic effects of systemic Formoterol at late stages after nerve injury and indicate the crucial role of microglial β2-AR in mediating the analgesic effects at early and late stages of mechanical allodynia and cold allodynia.

### Contribution of microglial *β*2-ARs to inhibitory effects of Formoterol on SNI-induced microgliosis and astrogliosis

Furthermore, we analyzed if deleting the microglial β2-AR influenced the accumulation of microglia in the SDH and the increased GFAP expression in activated astrocytes that are typical for SNI after six and 21 days post-nerve injury. Indeed, in contrast to control mice, Formoterol treatment failed to decrease the density of microglia in the SDH of *Cx3cr1-Adrb2^−/−^* mice of both sexes six and 21 days after SNI operation (Figure 7A, B), further supporting a role of β2-AR activation in the modulation of microglial reactivity and the induction of neuropathic pain.

**Figure 7.**
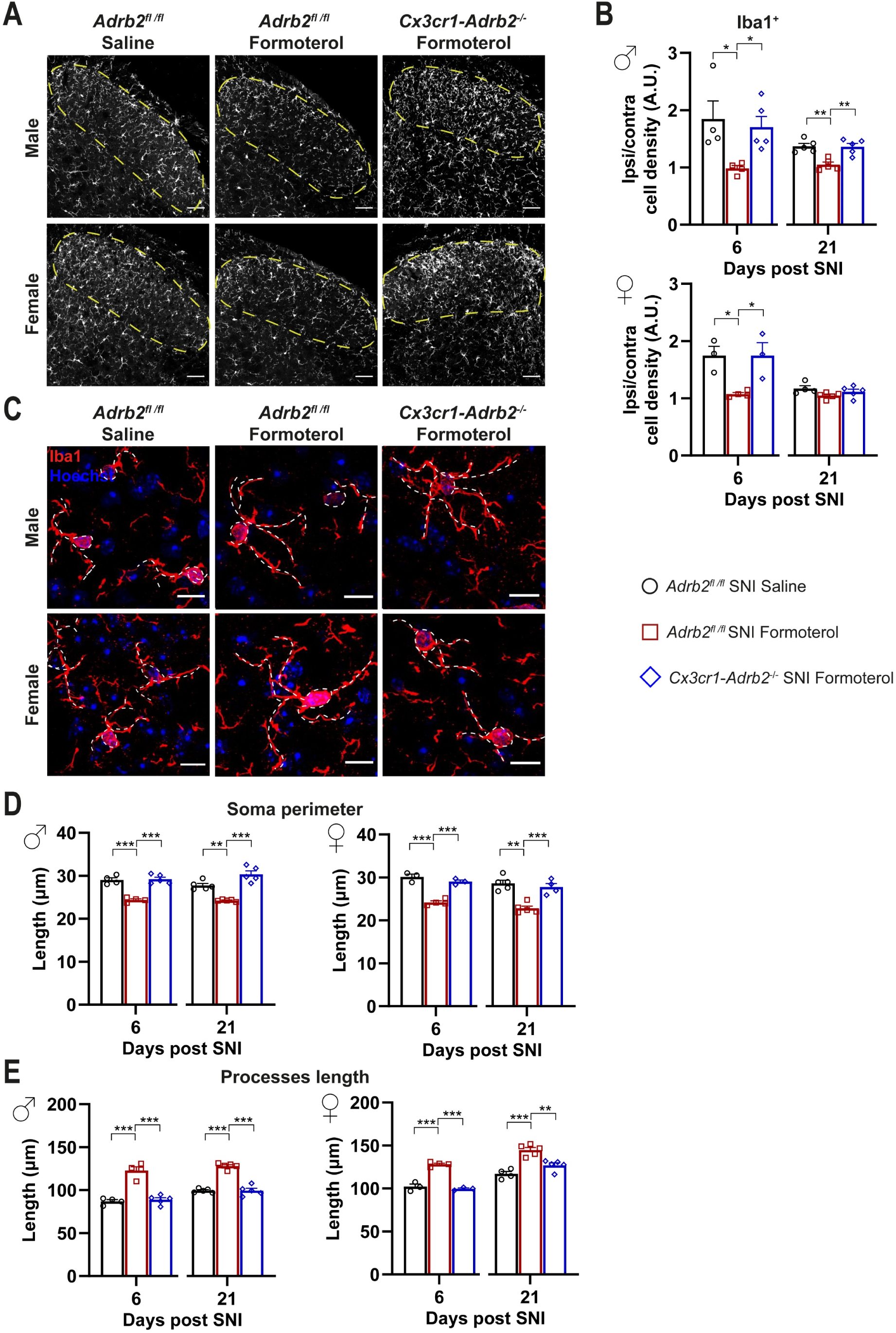
Spinal microglia response to the β2-AR agonist in SNI-operated control and transgenic mice. (A) Typical examples of Iba1-positive staining in the spinal dorsal horn (SDH) six days post nerve injury in control and transgenic mice who received saline or Formoterol injection. Scale bar = 60 μm. (B) Representative examples of microglia in the ipsilateral SDH in male and female mice, with somata and processes marked by white, dashed lines, six days after SNI operation injected with saline or Formoterol. Scale bar = 10 μm. (C) Quantitative analysis of microglia density (Iba1-positive cells) of control and *Cx3cr1-Adrb2^−/−^* male (left) and female (right) mice treated with saline, or Formoterol, six and 21 days after SNI operation. Ipsi/contra = ratio between the ipsilateral and contralateral SDH. A.U. = arbitrary unit. (D – E) Formoterol and saline application to control and transgenic male (left) and female (right) mice six or 21 days after SNI influences microglial morphological parameters: the perimeter of the soma (D) and the processes length (E). n = 3 – 5 / group; ordinary two-way ANOVA with main effects only was performed; * p < 0.05, ** p < 0.01, *** p < 0.001. Data are shown as mean ± SEM, individual data points are indicated.

In addition, Formoterol application failed to revert SNI-dependent morphological changes in *Cx3cr1-Adrb2*^−/−^ mice (Figure 7C, D), underlining the crucial role of microglial β2-AR for the analgesic effect of Formoterol. To confirm the role of microglial β2-AR in mediating the effect of Formoterol, we checked the expression of microglial activity markers (p-p38 and p-JNK) in mice that lost β2-AR receptors. As expected, contrary to what happened in control mice, in both male and female *Cx3cr1-Adr*b2^−/−^ mice Formoterol failed to damp the upregulation of microglial activity marker typically seen six and 21 days after SNI (Figure 8).

**Figure 8.**
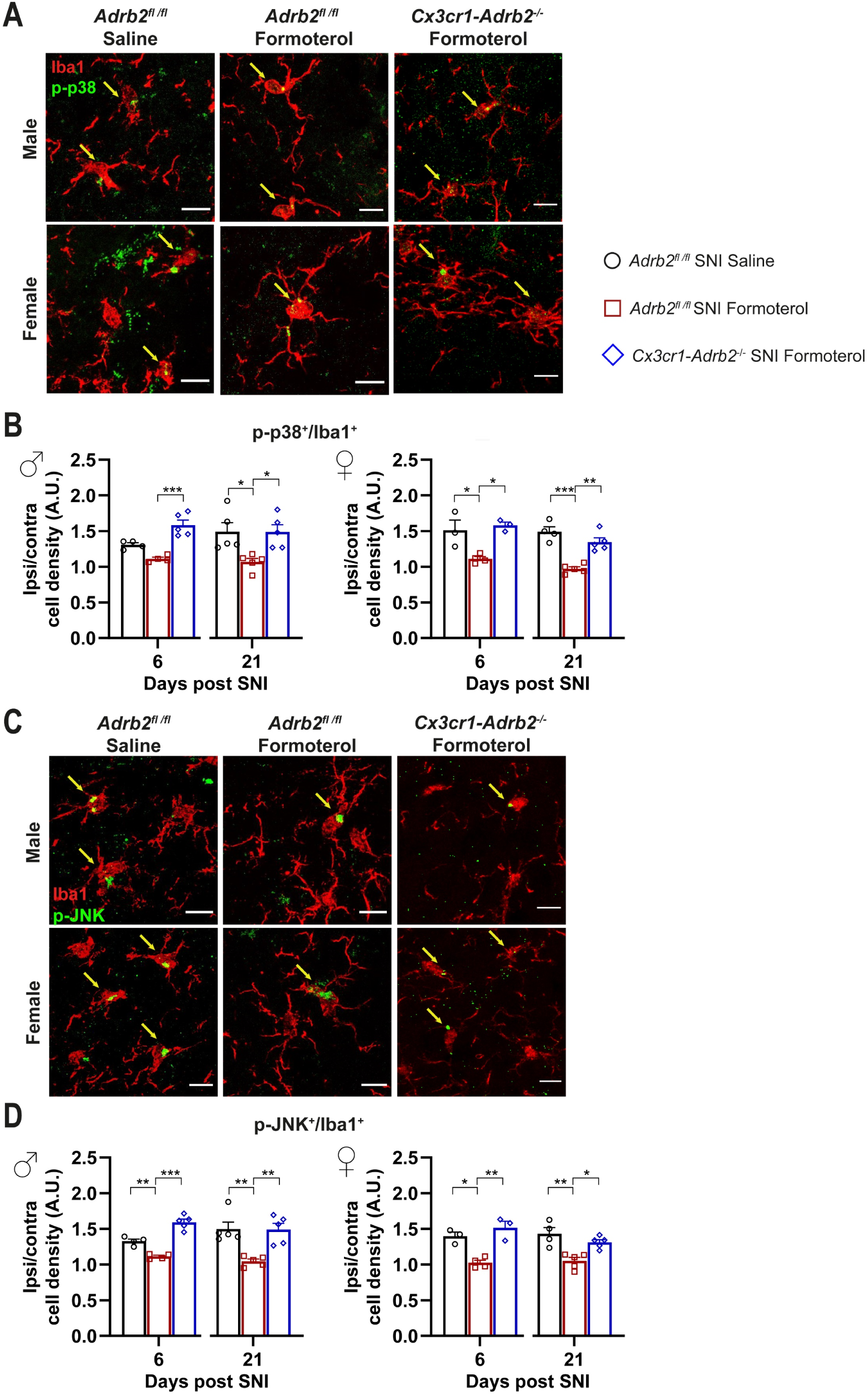
Microglial β2-AR knockdown impairs the Formoterol effect of lowering microglial activation markers levels in the spinal dorsal horn of SNI-operated mice. (A – B) Typical examples of colocalization of Iba1-positive cells and the microglial activation markers, p-p38 (A) and p-JNK (B) in the spinal dorsal horn (SDH) in male and female mice, six days after SNI operation in control and transgenic mice injected with saline or Formoterol. Double-positive cells are pointed by arrows. Scale bar = 10 μm. (C – D) Analysis of the co-staining of Iba-1 immunohistochemistry and the activation marker p-p38 (C) and Iba-1 positive cells and the activation marker p-JNK of control and *Cx3cr1-Adrb2^−/−^* male (left) and female (right) mice treated with saline or Formoterol, six and 21 days after SNI operation. Ipsi/contra = ratio between the ipsilateral and contralateral dorsal horn of the spinal cord. A.U. = arbitrary unit. n = 3 – 5 / group; ordinary two-way ANOVA with main effects only was performed; * p < 0.05, ** p < 0.01, *** p < 0.001. Data are shown as mean ± SEM, individual values are also displayed.

As with control mice, in *Cx3cr1-Adrb2^−/−^* of both sexes, we observed a significant increase in GFAP signal intensity at late time points (three weeks post-SNI) when neuropathic pain is fully established, but not over early time points (Figure 9A, B), which was partially, but significantly, inhibited by Formoterol (Figure 9A, B). Interestingly, also the nerve injury-induced upregulation of p-JNK in GFAP-positive cells in *Cx3cr1-Adrb2^−/−^* female mice at six and 21 days post-SNI was reversed by Formoterol (Figure 9C, D). Neither p-JNK upregulation nor effects of Formoterol were reliably seen in male and female mice at early time points after SNI (Figure 9D). This data indicates that at late time points after SNI operation, the contribution of the microglial β2-ARs to the effect of Formoterol is minimal.

**Figure 9.**
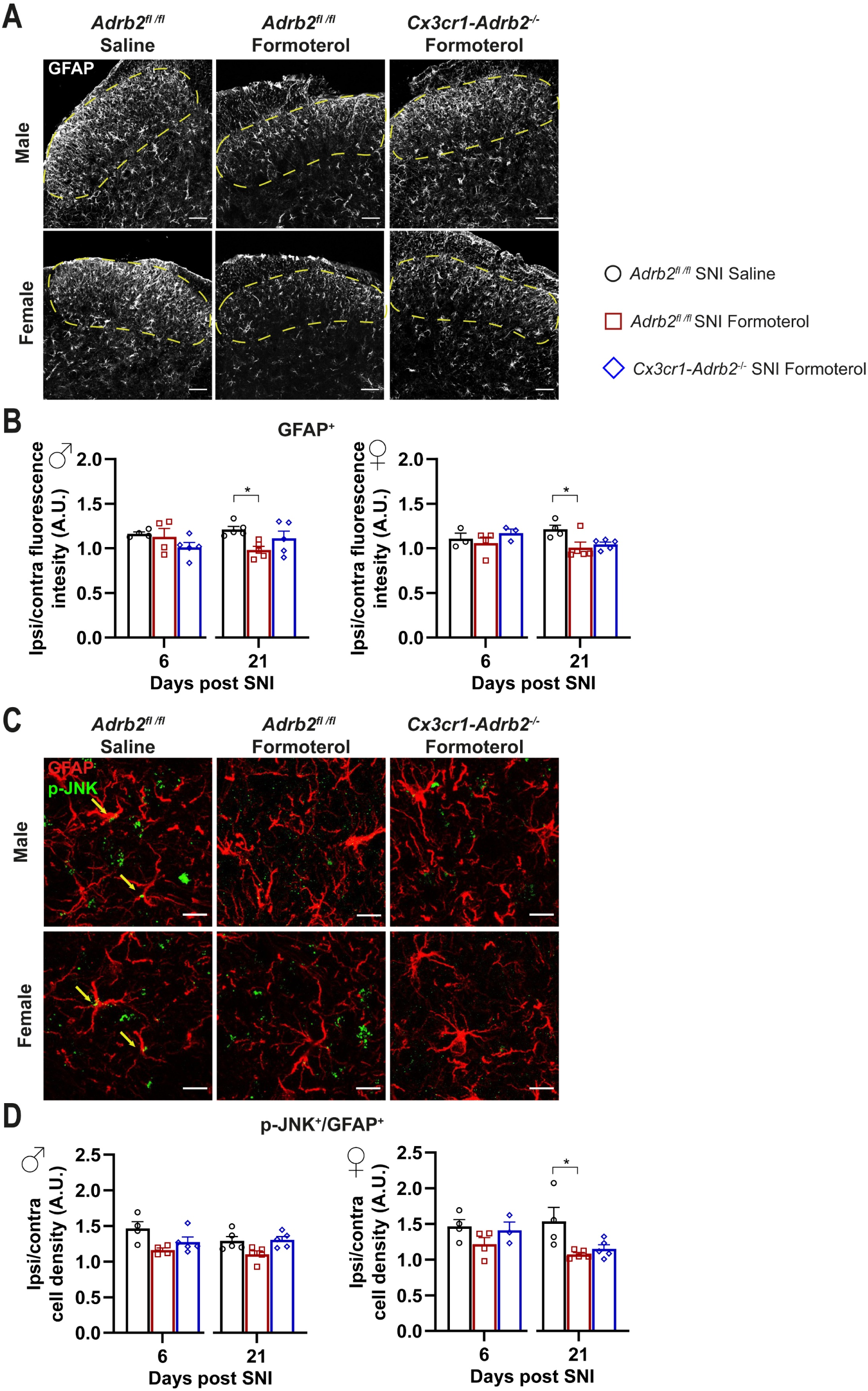
The knockdown of the microglial β2-AR did not affect the astrocytic response to Formoterol. (A) Examples of GFAP immune reactivity in the spinal dorsal horn 21 days after nerve injury in control and transgenic mice who received saline or Formoterol injection. Scale bar = 60 μm. (B) Representative examples of colocalization of GFAP astrocytic marker with glial activation marker, p- JNK in the spinal dorsal horn (SDH) of male and female mice, 21 days after SNI operation in mice injected with saline or Formoterol. Scale bar = 10 μm. (C) Analysis of GFAP fluorescent intensity of control and *Cx3cr1-Adrb2^−/−^* mice treated with saline, or Formoterol, six and 21 days after SNI operation. Ipsi/contra fluorescent intensity = ratio between values of fluorescent intensity obtained from the ipsilateral and contralateral SDH. (D) Quantitative analysis of the colocalization of GFAP and the activation marker p-JNK in the SDH of male (left) and female (right) mice treated with saline, or Formoterol, six and 21 days after SNI operation. Ipsi/contra = ratio between the ipsilateral and contralateral SDH. A.U. = arbitrary unit. n = 3 – 5 / group; ordinary two-way ANOVA with main effects only was performed. * p < 0.05. Data are shown as mean ± SEM, individual data points are given.

## Discussion

New therapies and deeper mechanistic insights are urgently needed for the clinical management of neuropathic pain. Noradrenergic signaling has long been known to play a key role in the pathophysiological sequelae leading to chronic pain, and drugs activating inhibitory (α2) noradrenergic receptors, such as clonidine, have been subject to intense clinical and preclinical investigation in the past [1, 41]. The present study now sets a focus on investigating glial noradrenergic signaling and its contributions via a mechanistically distinct class of receptors, namely excitatory receptors of the β2-type.

A few studies have reported beneficial effects of β2-AR agonists, such as terbutaline, on diverse types of inflammatory pain and more recently, also pain of neuropathic origin [42, 43]. In these studies, the authors show that, in the context of neuropathic pain, peripheral δ opioid (DOP) receptors expressed in Nav1.8+ neurons are required for the analgesic effect of β2-AR agonist Formoterol. However, the mechanistic link between these two receptor systems, opioids and adrenergic, is yet to be explored as well as the precise mode of action and the stage at which β2-AR activation would exhibit the best therapeutic effects on neuropathic pain remains unclear.

Here, we studied the impact of the pharmacological activation of β2-ARs on glial cells on the distinct phases of development and maintenance of neuropathic pain in the SNI model of neuropathic pain. This study goes beyond confirming previous positive reports on β2-ARs and reveals several novel insights, including: (i) β2-ARs expressed by microglia are required for the anti-allodynic effects of β2-AR agonists; (ii) β2-AR activation modulates different aspects and modalities of neuropathic pain in a stage-specific manner; (iii) β2-AR-mediated anti-allodynic actions at different temporal phases of neuropathic pain differentially involve microglia and astrocytes, and (iv) β2-AR activation not only suppresses the sensory component of neuropathic pain but also reduces negative affect at a time when neuropathic pain is chronically established.

Several observations support the view that β2-AR agonists exert analgesic actions in neuropathic pain primarily via microglia, at least over the period from early establishment of neuropathic hypersensitivity. One, we found that Formoterol induces strong molecular alterations in pure populations of microglia that promote an anti-inflammatory phenotype. Secondly, nerve injury-induced alterations in native populations of microglia in spinal dorsal horn circuits were fully reversed by Formoterol. Formoterol-induced suppression of microgliosis did not occur in mice with a microglia-specific loss of β2-ARs, showing that modulation of microglial function by Formoterol occurs via direct actions on β2-ARs in microglia and not downstream of β2-AR actions in other cell types that engage in paracrine signaling with microglia. Remarkably, Formoterol-induced reversal of neuropathic pain-associated behaviors was fully abrogated in mice with a microglia-specific loss of β2-ARs in both male and female mice. These findings suggest that over the initial phase post-injury when neuropathic pain develops, pro-inflammatory spinal cord microglial signaling can be hindered by the therapeutic use of β-AR agonists to put brakes on the sequence of pathophysiological events that lead to the full manifestation of neuropathic pain.

β2-ARs are expressed on the astrocytes, and indeed we observed inhibition of astrocyte activation by Formoterol in nerve-injured mice at later stages when neuropathic pain was chronically established in female mice. Although, there is evidence of sequential activation of microglia early post-injury and astrocytes thereafter, and there are strong indications of crosstalk between microglia and astrocytes. Our data indicate that β2-AR signaling is not involved in this crosstalk and that noradrenergic signaling in microglia and astrocytes occurs independently of each other. Our finding that Formoterol is efficacious in reducing allodynia when given at a late stage alone, and that this late effect is not mediated by microglial β2-AR implies that there are at least two, temporally distinct modes of action for the anti-allodynic effects of Formoterol: an early effect via microglia and a late effect via astrocytes. This implies that when patients seek therapy at late stages post-nerve injury when neuropathic pain has already become chronic, Formoterol can still be efficacious by acting via β2-AR in astrocytes. It must be acknowledged, however, that this study was limited to microglial manipulations and further studies involving astrocyte-specific deletion of *Adbr2* will further elucidate the role of astrocytes on the development and maintenance of chronic pain.

SNRIs act by increasing NA availability for signaling in neural pathways our results suggest that SNRIs-induced analgesia is likely to involve β2-AR signaling in microglia [44]. Our observation that β2-AR expression increases in microglia after nerve injury is supportive of this hypothesis. Furthermore, since astrocytes and microglia express a variety of adrenergic receptors [8, 9], and astrocytes are particularly rich in α2a-ARs [10], and our unpublished results], the contributions of these receptors remain to be clarified. Interestingly, the expression pattern of adrenergic receptors is dynamically regulated depending on the physiological or pathological state of neurons and glia. For example, treatment of microglia with lipopolysaccharides (LPS) leads to a decrease in the expression of β2-AR and upregulation of α2a-AR, which has a higher affinity for NA [9, 25]. It will be therefore of value in future experiments to study the dynamic interplay between the expression and function of diverse noradrenergic receptors in chronic pain conditions and utilize these data to design novel receptor-specific therapies that could be used complementary to SNRIs. Finally, only at 21 days after nerve injury, we observed sexual dimorphism in the action of microglia and astrocytes, and effects of β2-AR activation at early and late stages also hold implications for the choice of drugs and the optimal time frame of their use in therapy of neuropathic pain. It is interesting to consider how microglia are affected by β2-AR activation while suppressing neuropathic allodynia. In various pathological models such as Alzheimer’s or Parkinson’s, the application of norepinephrine has been shown to have an anti-inflammatory effect, while the blockade of β2-AR receptors by β-blockers promotes inflammation [45]. Furthermore, β2-AR agonists have been shown to suppress inflammatory phenotypic changes in activated macrophages and microglia through activation of the cyclic AMP-protein kinase A (cAMP-PKA) pathway, suppression of the production of pro-inflammatory molecules such as TNFα and IL-1β [46, 47], and by promoting the release of anti-inflammatory substances [48, 49]. β2-AR agonists were also reported to promote the conversion of LPS-activated microglia from an M1- to M2-like phenotype via the classical cAMP/PKA/cAMP response element-binding protein (CREB) pathway, as well as via phosphoinositide 3-kinase (PI3K) and p38 MAPK signaling [50]. Similarly, several other studies report that modulating the astrocytic β2-AR tone alters nuclear factor kappa-light-chain-enhancer of activated B cells (NF-κB)-dependent effects and the immune cell content of the CNS in pro-inflammatory conditions [51]. Taken together, the anti-allodynic actions of β2-AR, acting via microglia and astrocytes, can be likely attributed to the ability to hinder inflammatory crosstalk between neurons, glia, and other immune cells in neuroinflammation after nerve injury.

Lately, the scientific community has shown vast interest in determining the influence of sex differences in the role of microglia in hypersensitivity [52, 53]. Experiments in female mice that targeted the P2X4-BDNF-TRKB pathway and activation of p38 mitogen-activated protein kinase (MAPK) were ineffective in reducing pain hypersensitivity [54, 55]. Nevertheless, numerous studies report no evident sexual dimorphism in the analgesic effect of microglial inhibitors, genetic knockout of microglial-selective molecules, or ablation of microglia in various nerve injury models [56–60]. Our results show that microgliosis (increased density, morphology shifting to an ameboid phenotype, and upregulation of activation markers) is equally present in the SDH of both male and female mice three and six days after SNI and that Formoterol markedly lessens this aberrant upregulation. Regarding sex-specific changes, we found only a small number of notable differences in our behavior experiments. In the CPP test for spontaneous pain, soon after nerve injury, SNI-operated female mice do not show a significant preference towards the Formoterol-associated chamber, but only a trend. This indicates that a few days after nerve injury, the β2-AR agonist analgesic effect in female mice is not as efficacious in reducing spontaneous pain as in male mice. The most striking difference appears to be the differential involvement of microglia 21 days after the operation. In contrast with male mice, microglia do not accumulate in the SDH of SNI-operated female mice similar to the sham-operated ones. Moreover, only in female mice at late time point, Formoterol treatment fails to reverse the SNI-induced microglial enlargement of the soma and the shrinking of the processes and does not reduce the p-JNK signal in microglia. We can speculate that in female mice the microglia neuroimmune response is attenuated from the third week post-SNI. This suggests that microglia three weeks after SNI return to the basal resting state as the sham condition, in which the Formoterol effect is attenuated.

Furthermore, previous studies showed sex-independent astroglial signaling in the spinal cord in neuropathic pain [18], whereas others showed sex-dependent astrocytic response/uptake of glutamate and response to fatty acids in response to a brain injury [61]. In our study, Formoterol administration decreases p-JNK levels in astrocytes in the SDH of SNI-operated female mice but failed in male mice.

Finally, the observations of this study and previous studies on sexual dimorphism in the involvement of microglia and astrocytes [18, 53] and the effects of β2-AR activation at early and late stages also hold implications for the development of more target-specific drugs and the choice of the optimal time frame of their use in therapy of neuropathic pain.

## Supplementary Materials

Supplementary 1: SNI does not affect *Adrb2* mRNA expression in the spinal dorsal horn. β2-AR agonist attenuates inflammatory cytokines release from activated primary microglia and modulates spared nerve injury-induced sensitization in a time-dependent manner. Supplementary 2: Formoterol attenuates mechanical hypersensitivity in SNI-operated mice. Temporal scheme of conditional place preference (CPP) and female mice CPP heatmaps. Supplementary 3: Response of SNI-operated mice to a single late injection of Formoterol. Supplementary 4: Formoterol effect on microglia in the spinal dorsal horn of sham-operated mice and negative controls for secondary antibodies.

## Author Contributions

This work was conceived by MS, AA, and ED. Data were collected and analyzed by ED and MS. Manuscript was written by MS, ED and AA. All authors read and approved the final manuscript.

## Institutional Review Board Statement

All experimental procedures were approved by the local governing body (Regier-ungspräsidium Karlsruhe, Germany, Ref. 35-9185.81/G-177/17 and 35-9185.81/G-274/19) and abided by German Law that regulates animal welfare and the protection of animals used for the scientific purpose (TierSchG, TierSchVersV).

## Data Availability Statement

The data presented in this study are openly available upon reasonable request.

## Acknowledgments

The authors are grateful to C. Gartner for secretarial assistance, D. Baumgartl-Ahlert, T. Roth, and L.S. Grams for excellent technical assistance, Prof. R. Kuner (Pharmacology Institute, University of Heidelberg) for intellectual input and the help in writing the initial draft. We thank Prof. Frank Kirchhoff (CIPMM, Homburg, Saarland, Germany) for the intellectual input and fruitful discussions. We further thank Prof. Frank Kirchhoff (CIPMM, Homburg, Saarland, Germany), Dr. Steffen Jung (The Weizmann Institute of Science, Rehovot, Israel, and Prof. Gerald Karsenty (Columbia University, New York, NY, USA) for the *Cx3cr1*-creERT2 and *Adrb2*^fl/fl^ mice donation, respectively. Graphical abstract created with BioRender.com. The authors gratefully acknowledge the Interdisciplinary Neurobehavioral Core Facility of Medical Faculty Heidelberg for assistance with behavioral experiments and the data storage service (SDS@hd).

## References

1. Finnerup, N.B.; Kuner, R.; Jensen, T.S. Neuropathic Pain: From Mechanisms to Treatment. Physiol Rev 2021, 101, 259–301, doi:10.1152/physrev.00045.2019.

2. Finnerup, N.B.; Attal, N.; Haroutounian, S.; McNicol, E.; Baron, R.; Dworkin, R.H.; Gilron, I.; Haanpää, M.; Hansson, P.; Jensen, T.S.;, et al. Pharmacotherapy for neuropathic pain in adults: a systematic review and meta-analysis. Lancet Neurol 2015, 14, 162–173, doi:10.1016/S1474-4422(14)70251-0.

3. Caraci, F.; Merlo, S.; Drago, F.; Caruso, G.; Parenti, C.; Sortino, M.A. Rescue of Noradrenergic System as a Novel Pharmacological Strategy in the Treatment of Chronic Pain: Focus on Microglia Activation. Front Pharmacol 2019, 10, 1024, doi:10.3389/fphar.2019.01024.

4. Bahari, Z.; Meftahi, G.H. Spinal alpha2 -adrenoceptors and neuropathic pain modulation; therapeutic target. Br J Pharmacol 2019, 176, 2366–2381, doi:10.1111/bph.14580.

5. Fricker, L.D.; Margolis, E.B.; Gomes, I.; Devi, L.A. Five Decades of Research on Opioid Peptides: Current Knowledge and Unanswered Questions. Mol Pharmacol 2020, 98, 96–108, doi:10.1124/mol.120.119388.

6. Pan, H.L.; Wu, Z.Z.; Zhou, H.Y.; Chen, S.R.; Zhang, H.M.; Li, D.P. Modulation of Pain Transmission by G Protein-Coupled Receptors. Pharmacology & therapeutics 2008, 117, 141–141, doi:10.1016/J.PHARMTHERA.2007.09.003.

7. Liu, B.; Eisenach, J.C. Hyperexcitability of axotomized and neighboring unaxotomized sensory neurons is reduced days after perineural clonidine at the site of injury. J Neurophysiol 2005, 94, 3159–3167, doi:10.1152/jn.00623.2005.

8. Zhang, X.; Wang, J.; Qian, W.; Zhao, J.; Sun, L.; Qian, Y.; Xiao, H. Dexmedetomidine inhibits tumor necrosis factor-alpha and interleukin 6 in lipopolysaccharide-stimulated astrocytes by suppression of c-Jun N-terminal kinases. Inflammation 2014, 37, 942–949, doi:10.1007/s10753-014-9814-4.

9. Ishii, Y.; Yamaizumi, A.; Kawakami, A.; Islam, A.; Choudhury, M.E.; Takahashi, H.; Yano, H.; Tanaka, J. Anti-inflammatory effects of noradrenaline on LPS-treated microglial cells: Suppression of NFκB nuclear translocation and subsequent STAT1 phosphorylation. Neurochem Int 2015, 90, 56–66, doi:10.1016/j.neuint.2015.07.010.

10. Kremer, M.; Salvat, E.; Muller, A.; Yalcin, I.; Barrot, M. Antidepressants and gabapentinoids in neuropathic pain: Mechanistic insights. Neuroscience 2016, 338, 183–206, doi:10.1016/j.neuroscience.2016.06.057.

11. Old, E.A.; Clark, A.K.; Malcangio, M. The role of glia in the spinal cord in neuropathic and inflammatory pain. Handb Exp Pharmacol 2015, 227, 145–170, doi:10.1007/978-3-662-46450-2_8.

12. Donnelly, C.R.; Andriessen, A.S.; Chen, G.; Wang, K.; Jiang, C.; Maixner, W.; Ji, R.R. Central Nervous System Targets: Glial Cell Mechanisms in Chronic Pain. Neurotherapeutics 2020, 17, 846–860, doi:10.1007/s13311-020-00905-7.

13. Taves, S.; Berta, T.; Chen, G.; Ji, R.R. Microglia and spinal cord synaptic plasticity in persistent pain. Neural Plast 2013, 2013, 753656, doi:10.1155/2013/753656.

14. Hald, A.; Nedergaard, S.; Hansen, R.R.; Ding, M.; Heegaard, A.M. Differential activation of spinal cord glial cells in murine models of neuropathic and cancer pain. Eur J Pain 2009, 13, 138–145, doi:10.1016/j.ejpain.2008.03.014.

15. Gwak, Y.S.; Kang, J.; Unabia, G.C.; Hulsebosch, C.E. Spatial and temporal activation of spinal glial cells: Role of gliopathy in central neuropathic pain following spinal cord injury in rats. Experimental Neurology 2012, 234, 362–372, doi:10.1016/J.EXPNEUROL.2011.10.010.

16. Nam, Y.; Kim, J.H.; Kim, J.H.; Jha, M.K.; Jung, J.Y.; Lee, M.G.; Choi, I.S.; Jang, I.S.; Lim, D.G.; Hwang, S.H.;, et al. Reversible Induction of Pain Hypersensitivity following Optogenetic Stimulation of Spinal Astrocytes. Cell Rep 2016, 17, 3049–3061, doi:10.1016/j.celrep.2016.11.043.

17. Guan, Z.; Kuhn, J.A.; Wang, X.; Colquitt, B.; Solorzano, C.; Vaman, S.; Guan, A.K.; Evans-Reinsch, Z.; Braz, J.; Devor, M.;, et al. Injured sensory neuron-derived CSF1 induces microglial proliferation and DAP12-dependent pain. Nat Neurosci 2016, 19, 94–101, doi:10.1038/nn.4189.

18. Chen, G.; Zhang, Y.Q.; Qadri, Y.J.; Serhan, C.N.; Ji, R.R. Microglia in Pain: Detrimental and Protective Roles in Pathogenesis and Resolution of Pain. Neuron 2018, 100, 1292–1311, doi:10.1016/j.neuron.2018.11.009.

19. Lubahn, C.L.; Lorton, D.; Schaller, J.A.; Sweeney, S.J.; Bellinger, D.L. Targeting alpha- and beta-Adrenergic Receptors Differentially Shifts Th1, Th2, and Inflammatory Cytokine Profiles in Immune Organs to Attenuate Adjuvant Arthritis. Front Immunol 2014, 5, 346, doi:10.3389/fimmu.2014.00346.

20. Uzkeser, H.; Cadirci, E.; Halici, Z.; Odabasoglu, F.; Polat, B.; Yuksel, T.N.; Ozaltin, S.; Atalay, F. Anti-inflammatory and antinociceptive effects of salbutamol on acute and chronic models of inflammation in rats: involvement of an antioxidant mechanism. Mediators Inflamm 2012, 2012, 438912, doi:10.1155/2012/438912.

21. Zhang, F.F.; Morioka, N.; Abe, H.; Fujii, S.; Miyauchi, K.; Nakamura, Y.; Hisaoka-Nakashima, K.; Nakata, Y. Stimulation of spinal dorsal horn β2-adrenergic receptor ameliorates neuropathic mechanical hypersensitivity through a reduction of phosphorylation of microglial p38 MAP kinase and astrocytic c-jun N-terminal kinase. Neurochem Int 2016, 101, 144–155, doi:10.1016/j.neuint.2016.11.004.

22. Arora, V.; Morado-Urbina, C.E.; Gwak, Y.S.; Parker, R.A.; Kittel, C.A.; Munoz-Islas, E.; Miguel Jimenez-Andrade, J.; Romero-Sandoval, E.A.; Eisenach, J.C.; Peters, C.M. Systemic administration of a beta2-adrenergic receptor agonist reduces mechanical allodynia and suppresses the immune response to surgery in a rat model of persistent post-incisional hypersensitivity. Mol Pain 2021, 17, 1744806921997206, doi:10.1177/1744806921997206.

23. Albertini, G.; Etienne, F.; Roumier, A. Regulation of microglia by neuromodulators: Modulations in major and minor modes. Neurosci Lett 2020, 733, 135000, doi:10.1016/j.neulet.2020.135000.

24. Zhang, Y.; Chen, K.; Sloan, S.A.; Bennett, M.L.; Scholze, A.R.; O’Keeffe, S.; Phatnani, H.P.; Guarnieri, P.; Caneda, C.; Ruderisch, N.;, et al. An RNA-sequencing transcriptome and splicing database of glia, neurons, and vascular cells of the cerebral cortex. J Neurosci 2014, 34, 11929–11947, doi:10.1523/JNEUROSCI.1860-14.2014.

25. Gyoneva, S.; Traynelis, S.F. Norepinephrine modulates the motility of resting and activated microglia via different adrenergic receptors. J Biol Chem 2013, 288, 15291–15302, doi:10.1074/jbc.M113.458901.

26. Mori, K.; Ozaki, E.; Zhang, B.; Yang, L.; Yokoyama, A.; Takeda, I.; Maeda, N.; Sakanaka, M.; Tanaka, J. Effects of norepinephrine on rat cultured microglial cells that express alpha1, alpha2, beta1 and beta2 adrenergic receptors. Neuropharmacology 2002, 43, 1026–1034, doi:10.1016/s0028-3908(02)00211-3.

27. Hinoi, E.; Gao, N.; Jung, D.Y.; Yadav, V.; Yoshizawa, T.; Myers, M.G.; Chua, S.C.; Kim, J.K.; Kaestner, K.H.; Karsenty, G. The sympathetic tone mediates leptin’s inhibition of insulin secretion by modulating osteocalcin bioactivity. J Cell Biol 2008, 183, 1235–1242, doi:10.1083/jcb.200809113.

28. Yona, S.; Kim, K.W.; Wolf, Y.; Mildner, A.; Varol, D.; Breker, M.; Strauss-Ayali, D.; Viukov, S.; Guilliams, M.; Misharin, A.;, et al. Fate mapping reveals origins and dynamics of monocytes and tissue macrophages under homeostasis. Immunity 2013, 38, 79–91, doi:10.1016/j.immuni.2012.12.001.

29. Decosterd, I.; Woolf, C.J. Spared nerve injury: an animal model of persistent peripheral neuropathic pain. Pain 2000, 87, 149–158, doi:10.1016/S0304-3959(00)00276-1.

30. Yalcin, I.; Tessier, L.H.; Petit-Demoulière, N.; Waltisperger, E.; Hein, L.; Freund-Mercier, M.J.; Barrot, M. Chronic treatment with agonists of beta(2)-adrenergic receptors in neuropathic pain. Exp Neurol 2010, 221, 115–121, doi:10.1016/j.expneurol.2009.10.008.

31. Dixon, W.J. Efficient analysis of experimental observations. Annu Rev Pharmacol Toxicol 1980, 20, 441–462, doi:10.1146/annurev.pa.20.040180.002301.

32. Nees, T.A.; Wang, N.; Adamek, P.; Verkest, C.; La Porta, C.; Schaefer, I.; Virnich, J.; Balkaya, S.; Prato, V.; Morelli, C.;, et al. The molecular mechanism and physiological role of silent nociceptor activation. bioRxiv 2022.

33. Huang, E.Y.; Liu, T.C.; Tao, P.L. Co-administration of dextromethorphan with morphine attenuates morphine rewarding effect and related dopamine releases at the nucleus accumbens. Naunyn Schmiedebergs Arch Pharmacol 2003, 368, 386–392, doi:10.1007/s00210-003-0803-7.

34. Schweizerhof, M.; Stösser, S.; Kurejova, M.; Njoo, C.; Gangadharan, V.; Agarwal, N.; Schmelz, M.; Bali, K.K.; Michalski, C.W.; Brugger, S.;, et al. Hematopoietic colony-stimulating factors mediate tumor-nerve interactions and bone cancer pain. Nat Med 2009, 15, 802–807, doi:10.1038/nm.1976.

35. Bohlen, C.J.; Bennett, F.C.; Tucker, A.F.; Collins, H.Y.; Mulinyawe, S.B.; Barres, B.A. Diverse Requirements for Microglial Survival, Specification, and Function Revealed by Defined-Medium Cultures. Neuron 2017, 94, 759–773.e758, doi:10.1016/j.neuron.2017.04.043.

36. Costigan, M.; Scholz, J.; Woolf, C.J. Neuropathic pain: a maladaptive response of the nervous system to damage. Annu Rev Neurosci 2009, 32, 1–32, doi:10.1146/annurev.neuro.051508.135531.

37. King, T.; Vera-Portocarrero, L.; Gutierrez, T.; Vanderah, T.W.; Dussor, G.; Lai, J.; Fields, H.L.; Porreca, F. Unmasking the tonic-aversive state in neuropathic pain. Nat Neurosci 2009, 12, 1364–1366, doi:10.1038/nn.2407.

38. Pitzer, C.; Kuner, R.; Tappe-Theodor, A. EXPRESS: Voluntary and evoked behavioral correlates in neuropathic pain states under different housing conditions. Mol Pain 2016, 12, doi:10.1177/1744806916656635.

39. Chen, Z.; Doyle, T.M.; Luongo, L.; Largent-Milnes, T.M.; Giancotti, L.A.; Kolar, G.; Squillace, S.; Boccella, S.; Walker, J.K.; Pendleton, A.;, et al. Sphingosine-1-phosphate receptor 1 activation in astrocytes contributes to neuropathic pain. Proc Natl Acad Sci U S A 2019, 116, 10557–10562, doi:10.1073/pnas.1820466116.

40. Ji, R.R.; Suter, M.R. p38 MAPK, microglial signaling, and neuropathic pain. Mol Pain 2007, 3, 33, doi:10.1186/1744-8069-3-33.

41. Kuner, R.; Flor, H. Structural plasticity and reorganisation in chronic pain. Nat Rev Neurosci 2016, 18, 20–30, doi:10.1038/nrn.2016.162.

42. Ceredig, R.A.; Pierre, F.; Doridot, S.; Alduntzin, U.; Hener, P.; Salvat, E.; Yalcin, I.; Gaveriaux-Ruff, C.; Barrot, M.; Massotte, D. Peripheral Delta Opioid Receptors Mediate Formoterol Anti-allodynic Effect in a Mouse Model of Neuropathic Pain. Front Mol Neurosci 2019, 12, 324, doi:10.3389/fnmol.2019.00324.

43. Kremer, M.; Megat, S.; Bohren, Y.; Wurtz, X.; Nexon, L.; Ceredig, R.A.; Doridot, S.; Massotte, D.; Salvat, E.; Yalcin, I.;, et al. Delta opioid receptors are essential to the antiallodynic action of Beta2-mimetics in a model of neuropathic pain. Mol Pain 2020, 16, 1744806920912931, doi:10.1177/1744806920912931.

44. Damo, E. Glial cells as target for antidepressants in neuropathic pain. Neuroforum 2022, 28, 85–94, doi:10.1515/nf-2021-0036.

45. Evans, A.K.; Ardestani, P.M.; Yi, B.; Park, H.H.; Lam, R.K.; Shamloo, M. Beta-adrenergic receptor antagonism is proinflammatory and exacerbates neuroinflammation in a mouse model of Alzheimer’s Disease. Neurobiol Dis 2020, 146, 105089, doi:10.1016/j.nbd.2020.105089.

46. Izeboud, C.A.; Monshouwer, M.; van Miert, A.S.; Witkamp, R.F. The beta-adrenoceptor agonist clenbuterol is a potent inhibitor of the LPS-induced production of TNF-alpha and IL-6 in vitro and in vivo. Inflamm Res 1999, 48, 497–502, doi:10.1007/s000110050493.

47. Keranen, T.; Hommo, T.; Moilanen, E.; Korhonen, R. beta2-receptor agonists salbutamol and terbutaline attenuated cytokine production by suppressing ERK pathway through cAMP in macrophages. Cytokine 2017, 94, 1–7, doi:10.1016/j.cyto.2016.07.016.

48. Agac, D.; Estrada, L.D.; Maples, R.; Hooper, L.V.; Farrar, J.D. The beta2-adrenergic receptor controls inflammation by driving rapid IL-10 secretion. Brain Behav Immun 2018, 74, 176–185, doi:10.1016/j.bbi.2018.09.004.

49. Keranen, T.; Hommo, T.; Hamalainen, M.; Moilanen, E.; Korhonen, R. Anti-Inflammatory Effects of beta2-Receptor Agonists Salbutamol and Terbutaline Are Mediated by MKP-1. PLoS One 2016, 11, e0148144, doi:10.1371/journal.pone.0148144.

50. Sharma, M.; Arbabzada, N.; Flood, P.M. Mechanism underlying beta2-AR agonist-mediated phenotypic conversion of LPS-activated microglial cells. J Neuroimmunol 2019, 332, 37–48, doi:10.1016/j.jneuroim.2019.03.017.

51. Laureys, G.; Clinckers, R.; Gerlo, S.; Spooren, A.; Wilczak, N.; Kooijman, R.; Smolders, I.; Michotte, Y.; De Keyser, J. Astrocytic beta(2)-adrenergic receptors: from physiology to pathology. Prog Neurobiol 2010, 91, 189–199, doi:10.1016/j.pneurobio.2010.01.011.

52. Midavaine, E.; Cote, J.; Marchand, S.; Sarret, P. Glial and neuroimmune cell choreography in sexually dimorphic pain signaling. Neurosci Biobehav Rev 2021, 125, 168–192, doi:10.1016/j.neubiorev.2021.01.023.

53. Mogil, J.S. Qualitative sex differences in pain processing: emerging evidence of a biased literature. Nat Rev Neurosci 2020, 21, 353–365, doi:10.1038/s41583-020-0310-6.

54. Taves, S.; Berta, T.; Liu, D.L.; Gan, S.; Chen, G.; Kim, Y.H.; Van de Ven, T.; Laufer, S.; Ji, R.R. Spinal inhibition of p38 MAP kinase reduces inflammatory and neuropathic pain in male but not female mice: Sex-dependent microglial signaling in the spinal cord. Brain Behav Immun 2016, 55, 70–81, doi:10.1016/j.bbi.2015.10.006.

55. Sorge, R.E.; Mapplebeck, J.C.; Rosen, S.; Beggs, S.; Taves, S.; Alexander, J.K.; Martin, L.J.; Austin, J.S.; Sotocinal, S.G.; Chen, D.;, et al. Different immune cells mediate mechanical pain hypersensitivity in male and female mice. Nat Neurosci 2015, 18, 1081–1083, doi:10.1038/nn.4053.

56. Batti, L.; Sundukova, M.; Murana, E.; Pimpinella, S.; De Castro Reis, F.; Pagani, F.; Wang, H.; Pellegrino, E.; Perlas, E.; Di Angelantonio, S.;, et al. TMEM16F Regulates Spinal Microglial Function in Neuropathic Pain States. Cell Rep 2016, 15, 2608–2615, doi:10.1016/j.celrep.2016.05.039.

57. Peng, J.; Gu, N.; Zhou, L.; U, B.E.; Murugan, M.; Gan, W.B.; Wu, L.J. Microglia and monocytes synergistically promote the transition from acute to chronic pain after nerve injury. Nat Commun 2016, 7, 12029, doi:10.1038/ncomms12029.

58. Gu, N.; Eyo, U.B.; Murugan, M.; Peng, J.; Matta, S.; Dong, H.; Wu, L.J. Microglial P2Y12 receptors regulate microglial activation and surveillance during neuropathic pain. Brain Behav Immun 2016, 55, 82–92, doi:10.1016/j.bbi.2015.11.007.

59. Staniland, A.A.; Clark, A.K.; Wodarski, R.; Sasso, O.; Maione, F.; D’Acquisto, F.; Malcangio, M. Reduced inflammatory and neuropathic pain and decreased spinal microglial response in fractalkine receptor (CX3CR1) knockout mice. J Neurochem 2010, 114, 1143–1157, doi:10.1111/j.1471-4159.2010.06837.x.

60. Barragan-Iglesias, P.; Pineda-Farias, J.B.; Cervantes-Duran, C.; Bravo-Hernandez, M.; Rocha-Gonzalez, H.I.; Murbartian, J.; Granados-Soto, V. Role of spinal P2Y6 and P2Y11 receptors in neuropathic pain in rats: possible involvement of glial cells. Mol Pain 2014, 10, 29, doi:10.1186/1744-8069-10-29.

61. Crespo-Castrillo, A.; Arevalo, M.A. Microglial and Astrocytic Function in Physiological and Pathological Conditions: Estrogenic Modulation. Int J Mol Sci 2020, 21, doi:10.3390/ijms21093219.

