## Supplementary Figures for "Activation of β2-adrenergic receptors in microglia alleviates neuropathic hypersensitivity in mice"

A

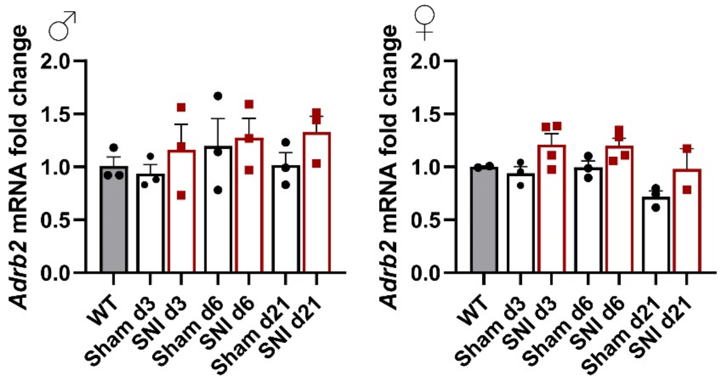


B

| | CSF-1 + vehicle | CSF-1 + Formoterol |
|---|---|---|
| **GM-CSF** | 1 ± 0.02 | 0.81 ± 0.03 |
| **INFγ** | 1 ± 0.02 | 0.43 ± 0.04 |
| **IL-1α** | 1 ± 0.01 | 0.86 ± 0.04 |
| **IL-9** | 1 ± 0.02 | 1.10 ± 0.02 |
| **IL-17** | 1 ± 0.01 | 0.82 ± 0.03 |
| **I-TAC** | 1 ± 0.01 | 0.84 ± 0.03 |
| **MIG** | 1 ± 0.01 | 1.12 ± 0.01 |
| **MIP-1γ** | 1 ± 0.03 | 1.29 ± 0.03 |
| **SDF-1** | 1 ± 0.02 | 0.80 ± 0.02 |
| **TCA-3** | 1 ±0.04 | 0.82 ± 0.03 |
| **TECK** | 1 ± 0.03 | 0.84 ± 0.02 |
| **TNF-α** | 1 ± 0.02 | 0.81 ± 0.03 |

C

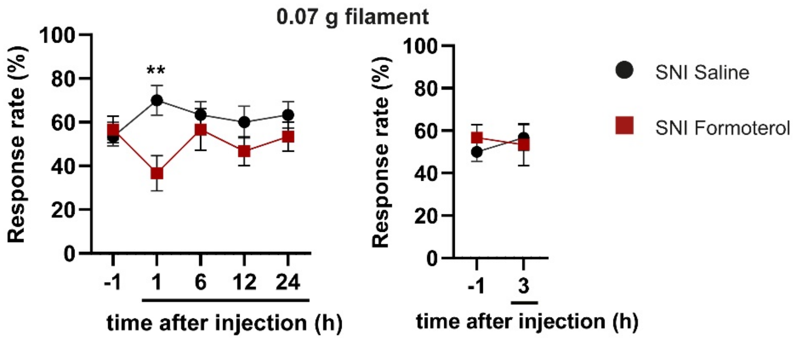


D

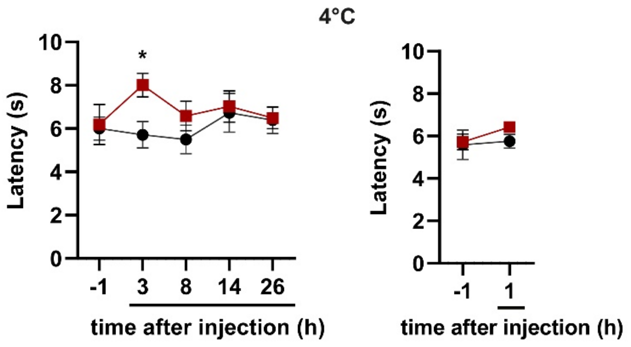

**Supplementary 1. SNI does not affect *Adrb2* mRNA expression in the spinal dorsal horn. β2-AR agonist attenuates inflammatory cytokines release from activated primary microglia and modulates spared nerve injury-induced sensitization in a time-dependent manner.** (A) Analysis of qPCR expression of *Adrb2* mRNA extracted from the spinal dorsal horn of male (left) or female mice (right) three, six, or 21 days after SNI or sham operation. n = 2 – 3. Data are expressed as mean ± SEM, individual values are also displayed. (B) Values of cytokines that significantly change in treated and untreated primary microglial cultures. (C – D) Time-course analysis of the effects of a single intraperitoneal (i.p.) dose of 50 µg/kg Formoterol on hind paw sensitivity to the 0.07 von Frey filament stimuli (C) or cold stimuli (D). Von Frey filaments test was applied on one group of SNI-operated mice 1, 6, 12, and 24 hours (h) after saline or Formoterol injection, whereas for a different group the mechanical hypersensitivity was measured 3 hours after the injection. Cold plate test was performed 3, 8, 14, and 26 h after injection for one group, but after 1 hour for a separate group. n = 6; two-tailed unpaired t-test was performed; * $p < 0.05$, ** $p < 0.01$, as compared between saline- and Formoterol-injected mice at the same time point.

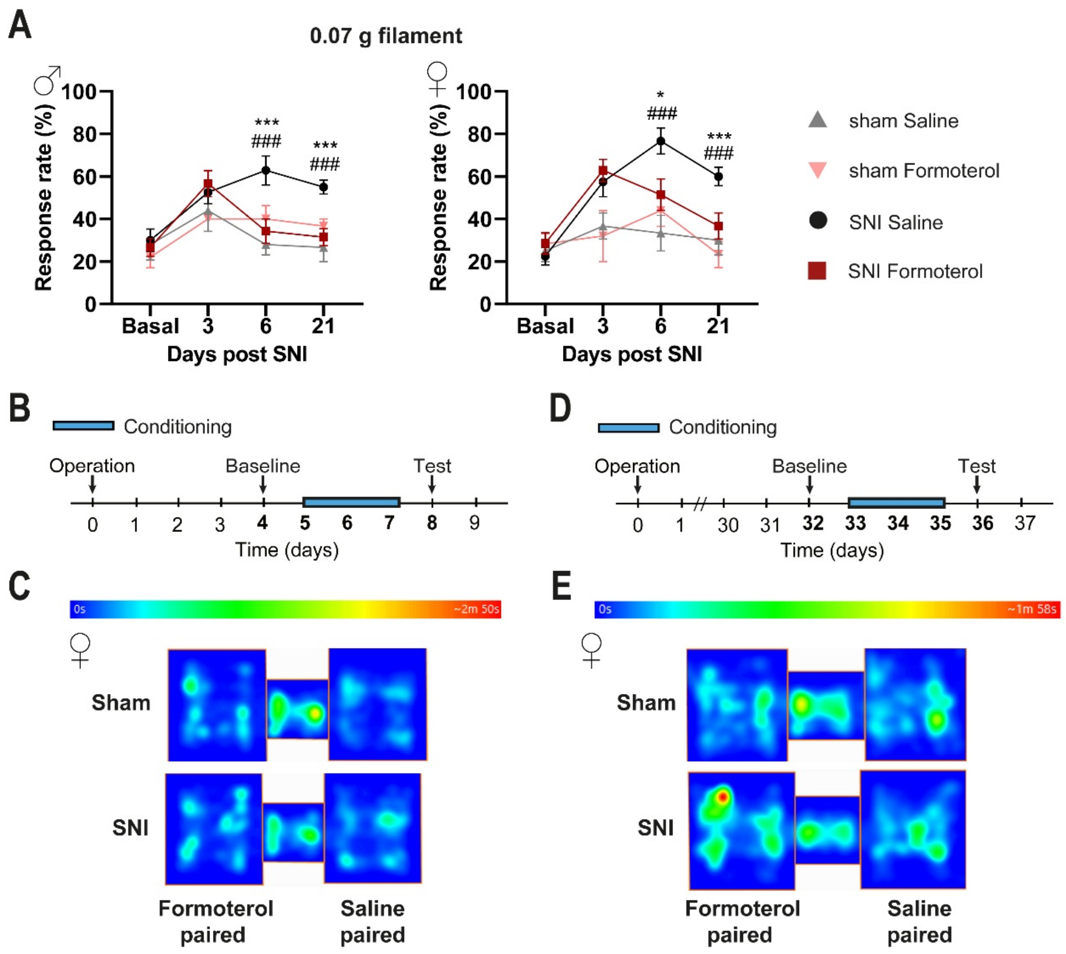


**Supplementary 2. Formoterol attenuates mechanical hypersensitivity in SNI-operated mice. Temporal scheme of conditional place preference (CPP) and female mice CPP heatmaps.**

(A) Mechanical sensitivity is displayed as response frequency to the 0.07 g filament in male (left) and female (right) before the SNI operation (basal measurement), three days after surgery, and six and 21 days after the SNI, one hour after Formoterol injection. n = 6 – 7 / group; t-test test was performed; * $p < 0.05$, *** $p < 0.001$ as compared SNI Saline and SNI Formoterol; ### $p < 0.001$ as compared SNI Saline and Sham Saline. (B - C) CPP for early time protocol after the operation (B) and example of SNI and sham female mice heat map tracking (C). (D - E) CPP for late time protocol after the surgery (C) and heat map example of SNI and sham female mice (E).

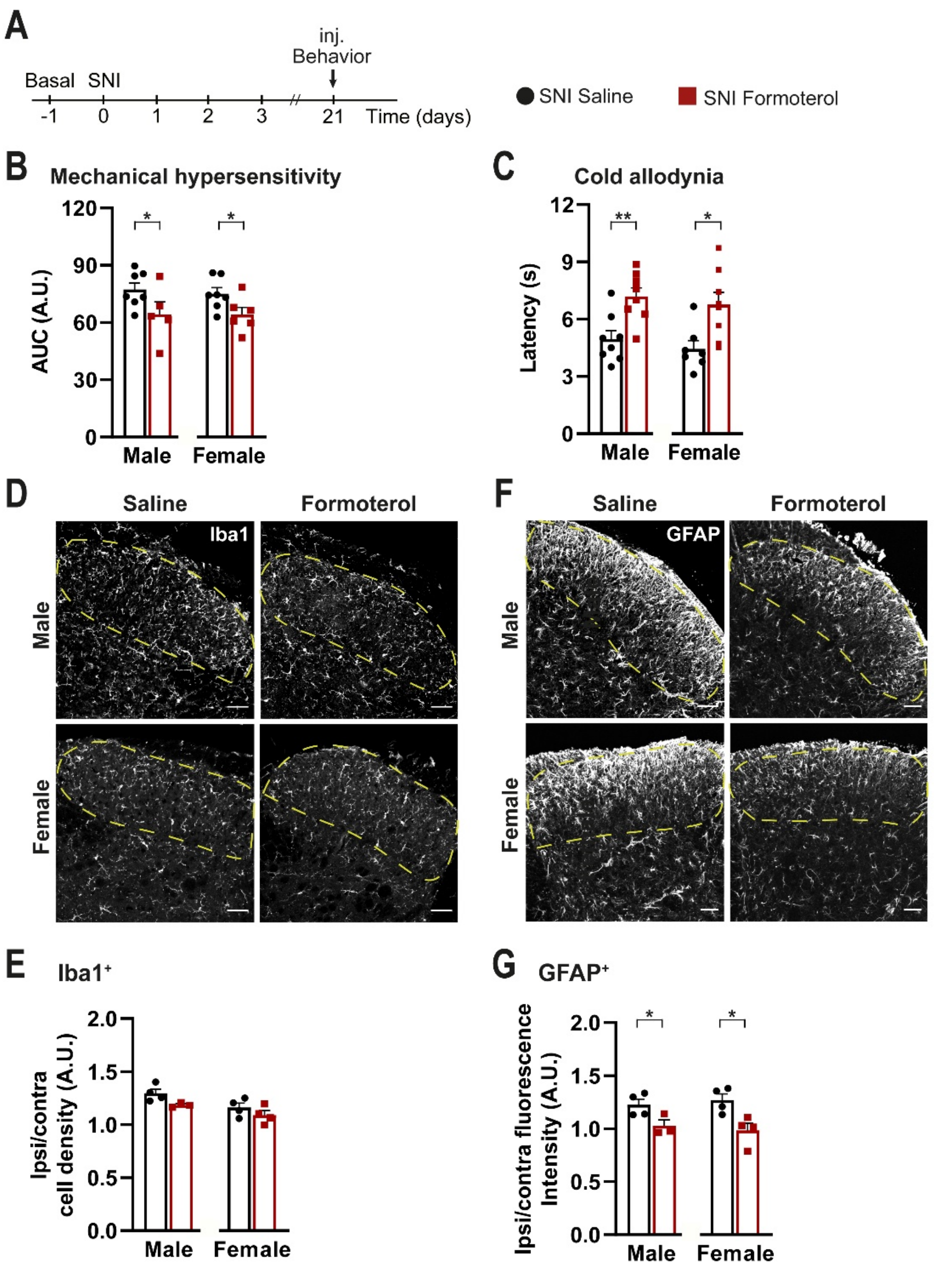


**Supplementary 3. Response of SNI-operated mice to a single late injection of Formoterol.** (A)Work plan for Von Frey and cold plate measurements on 8 weeks old mice. Inj. = injection. (B – C) Behavioral analysis of the effects of a single intraperitoneal injection of Formoterol on mechanical sensitivity displayed as integral of response frequency – von Frey force intensity (0.008 to 0.1 g) curves (AUC, A.U. = arbitrary unit) (B) and cold allodynia (C) given 21 days after SNI operation to mice of both genders. n = 5 – 8 / group. (D – G) Examples of Iba1 (D) or GFAP staining (F) and analysis of microglia density (E) or GFAP fluorescent intensity (G) in the spinal dorsal horn of male and female mice treated with one single injection of saline or Formoterol 21 days after SNI operation. Ipsi/contra = ratio between the ipsilateral and contralateral dorsal horn of the spinal cord. Ipsi/contra fluorescent intensity = ratio between values of fluorescent intensity obtained from the ipsilateral and contralateral dorsal horn of the spinal cord. Scale bar = 60 μm. n = 3 – 4 / group; two-tailed unpaired t-test was performed. * $p < 0.05$, ** $p < 0.01$. Data are expressed as mean ± SEM. Individual data points are given.

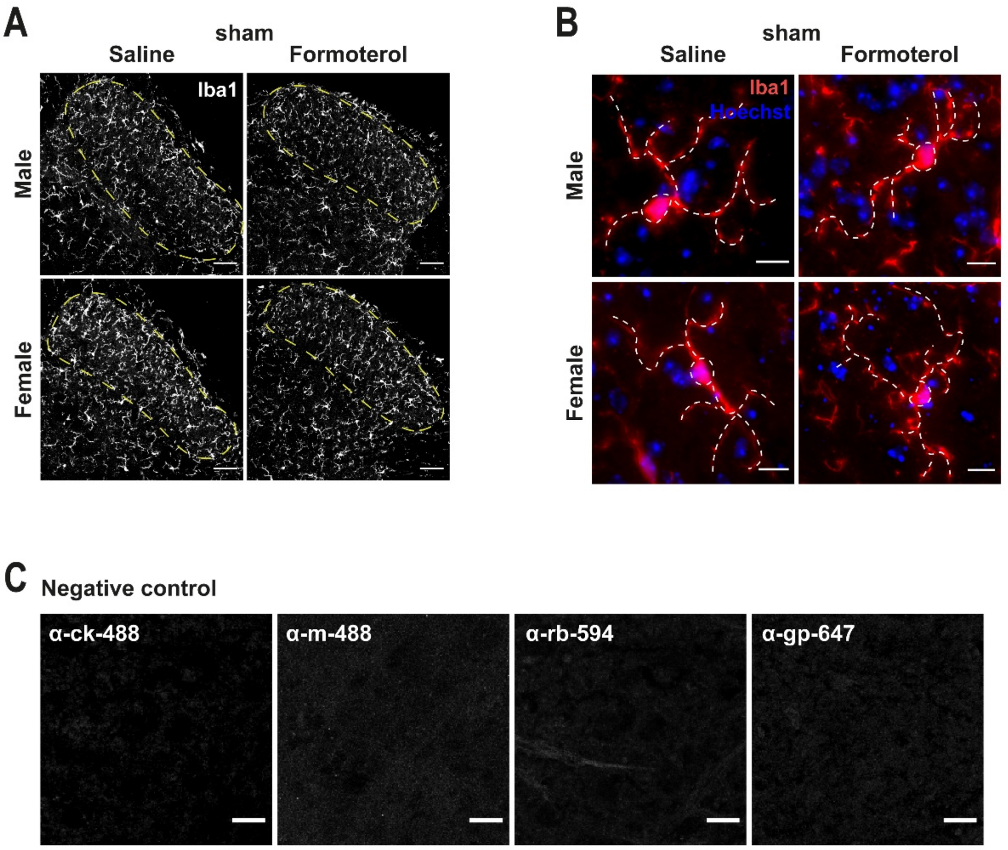


**Supplementary 4. Formoterol effect on microglia in the spinal dorsal horn of sham-operated mice and negative controls for secondary antibodies.**

(A) Typical examples of Iba1 positive cells in the ipsilateral spinal dorsal horn of male or female mice six days after the sham surgery, injected with saline or Formoterol. Scale = 60 μm. (B) Representative examples of microglia in the ipsilateral spinal dorsal horn in male and female mice, with somata and processes marked by white, dashed lines, six days after sham operation injected with saline or Formoterol. Scale bar = 10 μm. (C) Negative control staining for non-specific binding of the secondary antibodies. Scale bar = 10 μm.
